# Modulating AP-1 enables CAR-T cells to establish an intratumoral PD-1^+^Tcf1^+^ stem-like reservoir and overcomes resistance to PD-1 axis blockade

**DOI:** 10.1101/2025.04.10.648245

**Authors:** Andrew J. Snyder, Carolyn Shasha, Tam Ho, Sarah Garrison, Amelia R. Wilhelm, Mitchell G. Kluesner, Sergio Ortiz, W. Sam Nutt, Emma Bingham, Shruti S. Bhise, Everett Fan, Victor Zepeda, Megha Sarvothama, Xiao Wang, Shobha Potluri, Annalyssa Long, Anna Elz, Scott Furlan, Evan W. Newell, Shivani Srivastava

## Abstract

PD-1^+^Tcf1^+^ stem-like cells are critical mediators of endogenous T cell responses to PD-1/PD-L1 blockade and are maintained by MHC-dependent interactions with professional antigen-presenting cells (**APCs**). Unlike conventional T cells, CAR-T cells are activated by intact antigen expressed on tumors, not by peptide/MHC expressed on APCs, restricting their activation to the hostile tumor microenvironment (**TME**) that may impair preservation of this critical stem-like subset. Indeed, in an autochthonous model of ROR1^+^ lung cancer that we developed, CAR-T cells targeting the tumor-associated antigen ROR1 were uniformly Tcf1^+^ prior to infusion but rapidly downregulated Tcf1 *in vivo* and became terminally exhausted, similar to observations in patients, resulting in faster attrition and no enhancement in response to PD-L1 blockade. We hypothesized that overexpression of AP-1 family transcription factors, which can regulate T cell exhaustion, could enable CAR-Ts to maintain this critical PD-1^+^Tcf1^+^ subset within tumors independently of APCs and sensitize them to PD-1/PD-L1 blockade. Overexpression of the AP-1 TF c-Jun, but not BATF, improved preservation of PD-1^+^Tcf1^+^ CAR-T cells within tumors in a cell-intrinsic manner that correlated with increased persistence deeper within tumors. Notably, c-Jun overexpression alone was insufficient to prevent CAR-T exhaustion in the lung TME, in contrast to prior work in xenograft models, with progressive CAR-T dysfunction correlated with PD-1-dependent downregulation of c-Jun. However, c-Jun overexpression dramatically sensitized CAR-Ts to PD-L1 blockade, which restored c-Jun levels in CAR-Ts, drove log-fold expansion of CAR-Ts within tumors, and induced nearly complete eradication of ROR1^+^ tumor in highly aggressive models of lung cancer. Altogether, our data show that combination with PD-L1 blockade is necessary to unleash the full potential of c-Jun-overexpressing CAR-T cells in aggressive solid tumors like lung cancer and suggest that strategies to enhance formation of intratumoral PD-1^+^Tcf1^+^ reservoirs can overcome CAR-T resistance to PD-1 blockade.

## INTRODUCTION

Immune checkpoint inhibitors (**ICI**) targeting the PD-1/PD-L1 axis have been transformative for the treatment of lung cancer, which is the leading cause of cancer mortality in the United States and worldwide,^1^ but ∼80% of lung cancer cases remain refractory to ICI.^2^ One therapeutic strategy is to infuse patients with T cells engineered to express chimeric antigen receptors (**CAR**) specific for tumor antigens, endowing patients with a supply of functional tumor-specific T cells. Immunotherapy using CAR-T cells targeting CD19 has induced dramatic remissions in some patients with advanced hematological malignancies, representing one of the most remarkable therapeutic advances in the past decade.^3–12^ Extending this success to more common epithelial cancers like lung cancer, however, has proved more challenging.^1,13^ Efficacy of CAR-T cell therapies in solid tumors has been limited by multiple factors, including heterogeneity of tumor antigen expression, T cell dysfunction in the tumor microenvironment (**TME**), and poor persistence of infused T cells.^13–18^ The tumor-associated antigen ROR1 was previously identified as an attractive target for CAR-T cells due to its high expression in multiple common, incurable cancers, including non-small cell lung cancer (**NSCLC**), triple-negative breast (**TNBC**), and chronic lymphocytic leukemia (**CLL**), and limited expression in vital adult tissues.^19–22^ In a phase 1 clinical trial conducted at our Center, CAR-T cells targeting ROR1 induced responses in CLL patients but upregulated multiple inhibitory receptors, became dysfunctional, and persisted poorly in TNBC and NSCLC patients, suggesting that strategies to preserve CAR-T function are needed to extend efficacy to more common solid tumors.^21,23^ An obvious strategy to rescue CAR-T cell function is to combine treatment with ICI; however, clinical studies thus far have shown mixed results as to whether ICI enhances CAR-T cell efficacy, and the mechanisms underlying CAR-T resistance to ICI are unclear.^24–34^ Understanding the pathways that limit T cell function in the context of PD-1/PD-L1 blockade, thus, is critical for developing more effective immunotherapies for patients with aggressive solid tumors.

Recent work has demonstrated that a subset of PD-1^+^Tcf1^+^ exhausted T cells, termed precursor exhausted cells (**Tpex**), are critical for sustaining immune responses to tumors and chronic infections.^35–43^ T cell exhaustion is a progressive decline in effector function and memory potential driven by chronic antigen stimulation that is commonly observed in cancer and chronic infection.^44–49^ PD-1^+^Tcf1^+^ Tpex retain some stem-like qualities, including the capacity for self-renewal, proliferation, and differentiation into transitory effector-like cells that mediate tumor cell killing.^35–43^ Tpex provide the proliferative burst after PD-1 blockade, and strategies to deplete or boost Tpex can shorten or extend tumor control and response to PD-1 blockade, respectively.^38,36,38,39,50,51^ Unlike terminally exhausted cells, which are scattered throughout tumors, Tpex preferentially co-localize with antigen-presenting cells (**APCs**) in tumor-draining lymph nodes (**tdLN**) and perivascular niches, where they are maintained by MHC-dependent interactions.^36,51–57^ This spatial segregation has been observed in patients and multiple animal models of cancer, chronic infection, and autoimmunity, suggesting that localization of Tpex to APC-rich regions may protect them from terminal differentiation.^36,42,53–55,58,59^ The signals provided in APC niches that promote Tpex maintenance are still being uncovered and may involve protection from more persistent antigen stimulation, inhibition of mTOR signaling, and/or reduced hypoxia, all of which are more prevalent in the TME and promote terminal exhaustion.^39,41,50,52,60^

Unlike conventional T cells, CAR-T cells are primarily activated through their CAR by intact antigen expressed on tumor cells, not by peptide/MHC complexes expressed on tumor cells and APCs. This MHC-independent design is commonly regarded as a positive trait, as it enables CAR-T cells to kill tumors that have downregulated antigen presentation on MHC, a common mechanism of tumor escape from MHC-restricted T cells.^61^ However, this design may also act as an Achilles’ heel for CAR-T cells, as it restricts their activation to the TME, a more hostile environment than APC niches that may promote Tcf1 downregulation and inhibit preservation of stem-like qualities. Given the critical role of APCs and tdLNs in regulating and maintaining Tpex, it is unclear how insufficient activation of CAR-T cells by APCs impacts their ability to establish and maintain an *in vivo* reservoir of stem-like PD-1^+^Tcf1^+^ cells and how this affects their response to PD-1 blockade.

Defining the mechanisms limiting CAR-T cell function and sensitivity to ICI in solid tumors requires more sophisticated and rigorous animal models than are routinely in use. We previously adapted the Kras^LSL-G12D/+^;p53^fl/fl^ (**KP**) autochthonous mouse model of NSCLC to express human ROR1 (**KP^ROR^**^1^) to evaluate strategies for enhancing the efficacy of ROR1 CAR-T cells.^21,62^ Infecting KP mice intratracheally with lentivirus expressing Cre recombinase both deletes p53 and activates oncogenic Kras^G12D^ in some lung epithelial cells, resulting in development of tumors that co-evolve naturally with the host immune system, develop a highly suppressive TME that resembles human NSCLC, and are refractory to ICI.^62–64^ We demonstrated that this aggressive model recapitulates the poor persistence and dysfunction of ROR1 CAR-T cells observed in patients.^21^ Moreover, tumors progress slowly over 4-5 months, enabling long-term analyses of CAR-T function and differentiation that are not possible in rapidly progressive transplantable and xenograft models.^64^ The KP^ROR1^ model, thus, closely mirrors observations in patients and provides a unique platform to test strategies to enhance CAR-T cell activity and sensitivity to ICI that can inform clinical translation.

An advantage of adoptive cell therapy is the opportunity to engineer tumor-reactive T cells to promote their maintenance of PD-1^+^Tcf1^+^ Tpex and preserve sensitivity to ICI. Exhaustion develops in part due to an imbalance between NFAT and activating AP-1 transcription factors (**TFs**), with insufficient activation of AP-1 TFs promoting “exhausted” rather than functional “effector” T cell differentiation.^45,65^ Because overexpression of AP-1 TFs have been shown to regulate T cell exhaustion,^65,66^ we hypothesized that modulation of AP-1 TFs could protect CAR-T cells from terminal differentiation and enable them to establish an *in vivo* reservoir of stem-like PD-1^+^Tcf1^+^ cells that could sensitize them to further enhancement with PD-1 blockade. We show that, although ROR1 CAR-T cells are uniformly Tcf1^+^ prior to infusion, they rapidly downregulate Tcf1 *in vivo*, fail to establish a PD-1^+^Tcf1^+^ reservoir, and undergo terminal exhaustion, similar to observations in patients. In the absence of a Tpex reservoir, PD-L1 blockade did not enhance ROR1 CAR-T activity and instead drove their terminal differentiation and more rapid attrition within tumors. Here, we use highly aggressive models of ROR1^+^ NSCLC to define the mechanisms underlying CAR-T resistance to PD-1 axis blockade and evaluate strategies to enhance formation of an intratumoral reservoir of PD-1^+^Tcf1^+^ stem-like CAR-T cells that can restore CAR-T cell sensitivity to PD-1 axis blockade.

## RESULTS

### ROR1 CAR-T cells fail to maintain a Tcf1+ reservoir within lung tumors and are not enhanced by PD-L1 blockade

We previously adapted the Kras/p53 autochthonous mouse model of lung adenocarcinoma to express ROR1 by modifying the lentivirus used to induce tumors to co-express Cre and truncated human ROR1 (**hROR1**).^21^ Because linkage of multiple transgenes via 2A ribosomal skip elements can result in decreased co-expression of promoter-distal transgenes,^67^ we modified the lentivirus used to induce tumors to place hROR1 in the 5’ position (**hROR1-P2A-Cre**), which increased co-expression of Cre and hROR1 without increasing hROR1 levels (Fig. S1A, B).^21^ In KP mice, hROR1-P2A-Cre lentivirus induced lung tumors that showed hROR1 expression in ∼50-70% of tumor cells by 12 weeks post-infection, when tumors were visible by microCT, and this fraction remained stable up to 18 weeks post-infection (Fig. S1C). Heterogeneous ROR1 expression could arise from silencing of the hROR1-P2A-Cre transgene or transient Cre expression without stable lentiviral integration in a subset of cells.^68^ Nevertheless, this level of heterogeneity is similar to that found in patients with ROR1^+^ TNBC and NSCLC, in which ROR1^+^ tumors express ROR1 in >50% of tumor cells.^19^ The KP^ROR1^ model, thus, enables us to study ROR1 CAR-T cell therapy directed against a clinically relevant antigen expressed at physiological levels.

To target hROR1, we used a previously described ROR1 CAR comprised of a single-chain variable fragment (**scFv**) that recognizes a similar epitope as the CAR used in the clinical trial and CD28 and CD3ζ signaling domains (Fig. 1A).^69,70^ When administered to tumor-bearing KP^ROR1^ mice after lymphodepleting chemotherapy with cyclophosphamide (**Cy**), ROR1 CAR-T cells accumulated within KP^ROR1^ tumors, upregulated the co-inhibitory receptors PD-1 and TIM-3, and lost the ability to produce IFNγ upon restimulation *ex vivo* compared to control T cells modified to express only the truncated murine CD19 transduction marker (**tCD19**) (Fig. 1A, 1B), indicating that ROR1 CAR-T cells undergo terminal exhaustion in the KP^ROR1^ model.^21^ Interestingly, ROR1 CAR-T cells prior to infusion highly expressed Tcf1, a TF critical for maintaining the stem-like features of memory and Tpex cells, and did not express PD-1, exhibiting a central memory-like phenotype that has been associated with improved clinical responses (Fig. 1C).^71–77^ However, upon infusion into tumor-bearing mice, ROR1 CAR-T cells rapidly and progressively downregulated Tcf1 and upregulated PD-1, adopting a PD-1^+^Tcf1^−^TIM-3^+^ phenotype that has been associated with poor response to PD-1 axis blockade.^78^ Indeed, even though ROR1 CAR-T cells highly expressed PD-1 and PD-L1 was highly upregulated on tumor neutrophils and macrophages in mice treated with ROR1 CAR-T cells (Fig. 1D), suggesting increased activation of the PD-1/PD-L1 pathway, PD-L1 blockade did not enhance ROR1 CAR-T cell activity. While PD-L1 blockade drove increased PD-1 expression and activation of ROR1 CAR-T cells, it did not increase numbers of ROR1 CAR-T cells and instead drove more rapid CAR-T cell attrition within KP^ROR1^ tumors (Fig. 1E). Consequently, combination treatment with anti-PD-L1 did not improve ROR1 CAR-T cell efficacy, with ROR1^+^ tumor progressing similarly in both groups (Fig. 1F, 1G). Thus, despite their high expression of Tcf1 pre-infusion, ROR1 CAR-T cells fail to establish a reservoir of PD-1^+^Tcf1^+^ cells *in vivo*, and this correlates with their terminal exhaustion and inability to be enhanced by PD-L1 blockade.

**Fig. 1.**
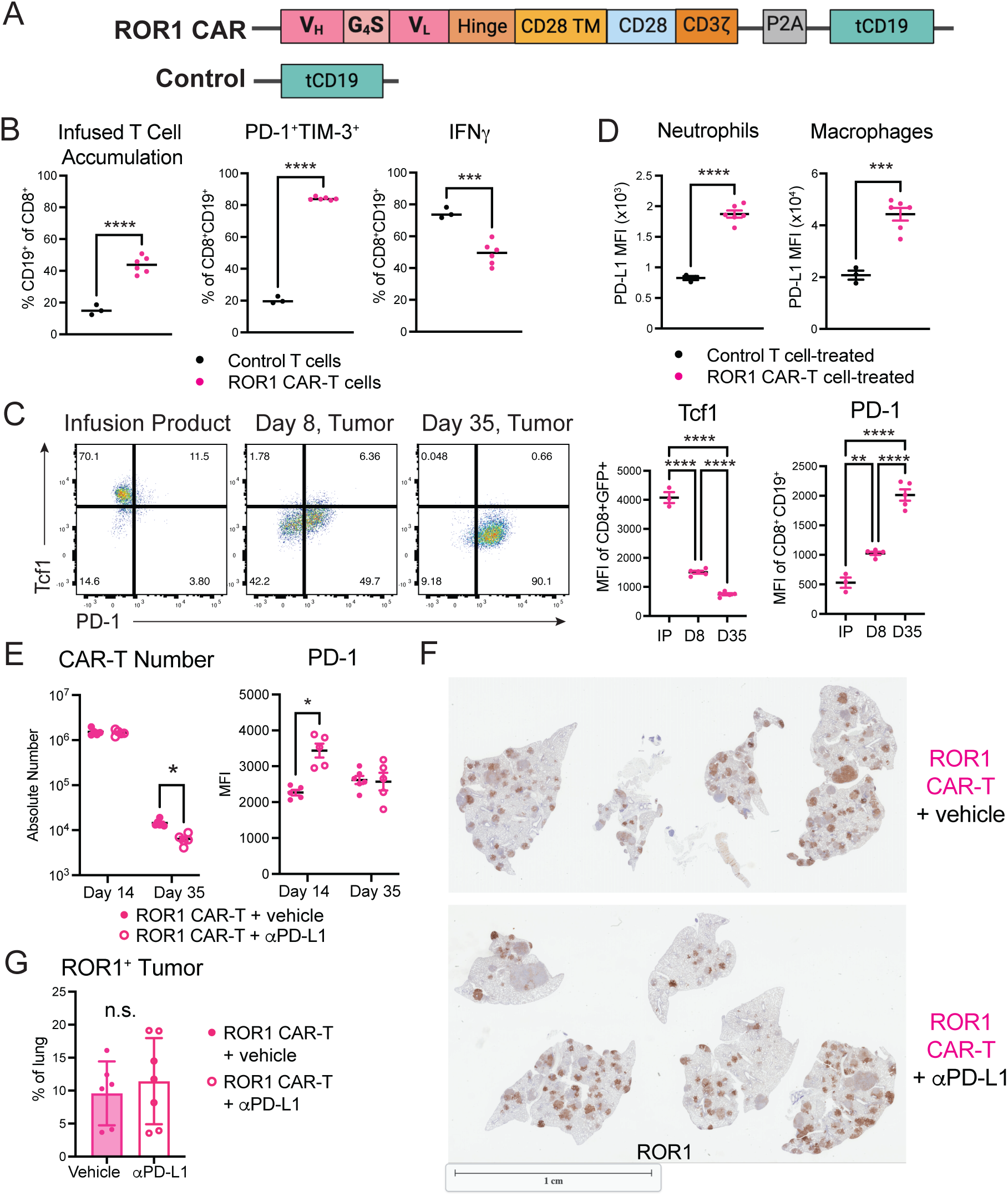
ROR1 CAR-T cells fail to maintain a Tcf1^+^ reservoir within lung tumors and are not enhanced by PD-L1 blockade. A) Schematic of retroviral constructs used to engineer murine ROR1 CAR-T and control T cells. TM = transmembrane. tCD19 = truncated murine CD19. B) Frequency (left), PD-1 and TIM-3 expression (middle), and IFNγ production upon *ex vivo* restimulation with PMA and ionomycin (right) by infused CD8^+^CD19^+^ T cells in lungs 8 days post-infusion into KP^ROR1^ mice. N=3-6 mice per group. Unpaired Student’s two-way t-test. C) Representative flow plots (left) and summary (right) of Tcf1 and PD-1 expression by CD8^+^CD19^+^ CAR-T cell over time *in vivo*. N=5 mice per group. One-way ANOVA with Tukey’s post-test. D) PD-L1 expression on CD11b^+^Ly6G^+^ neutrophils and CD11c^+^F4/80^+^ macrophages within KP^ROR1^ lung tumors 8 days post-treatment with control T cells or ROR1 CAR-T cells. N=3-6 mice per group. Unpaired Student’s two-way t-test. E) ROR1 CAR-T cell number (left) and PD-1 expression (right) 14 and 35 days post-infusion into KP^ROR1^ mice and treatment with vehicle or anti-PD-L1. N=5-6 mice per group. Unpaired Student’s two-way t-test. F) Representative IHC staining for ROR1 on lungs of KP^ROR1^ mice 45 days post-infusion with ROR1 CAR-T cells +/− anti-PD-L1. G) Quantification of ROR1^+^ tumor from IHC images of KP^ROR1^ lungs treated as indicated 45 days post-infusion. N=6-7 mice per group. Unpaired Student’s two-way t-test. Data are representative of 2 independent experiments.

### Overexpression of AP-1 family TFs affects maintenance of intratumoral Tcf1^+^ CAR-T cells and long-term persistence in vivo

We hypothesized that strategies to enhance preservation of PD-1^+^Tcf1^+^ CAR-T cells *in vivo* could both improve CAR-T cell persistence and sensitize them to further enhancement with ICI. T cell exhaustion is driven by a balance between NFAT and AP-1, and overexpression of AP-1 family TFs like c-Jun and BATF have been shown to partially restore CAR-T cell function in transplantable and xenograft tumor models.^65,66^ To examine how overexpression of these AP-1 family TFs affected the ability of CAR-T cells to establish a PD-1^+^Tcf1^+^ reservoir within tumors, we modified the CAR construct to co-express murine c-Jun or BATF upstream of the ROR1 CAR and a tCD19/GFP transduction marker (**cJun.ROR1** and **BATF.ROR1 CAR-T cells,** respectively) (Fig. 2A). Prior to infusion, cJun.ROR1 and BATF.ROR1 CAR-T cells expressed comparable levels of surface CAR and higher levels of c-Jun and BATF, respectively, compared to ROR1 CAR-T cells (Fig. S2A). Additionally, cJun.ROR1 and BATF.ROR1 CAR-T cells showed similarly high Tcf1 expression, proliferative capacity, cytokine production, and cytotoxic degranulation upon stimulation as ROR1 CAR-T cells prior to infusion, though their capacity to produce IL-2 was slightly increased (Fig. S2B-E). These data suggest that c-Jun or BATF overexpression do not substantially alter CAR-T functional capacity prior to infusion or upon acute stimulation *in vitro*. To determine how overexpression of c-Jun or BATF affected maintenance of Tcf1^+^ CAR-T cells *in vivo*, we transplanted B6 mice with lung tumors using a KP-ROR1 cell line derived from the GEM model and treated them with Cy and ROR1 CAR-T, cJun.ROR1 CAR-T, or BATF.ROR1 CAR-T cells. Whereas tumor-infiltrating ROR1 CAR-T cells and BATF.ROR1 CAR-T cells rapidly downregulated Tcf1 by 8 days post-infusion, cJun.ROR1 CAR-T cells maintained significantly higher frequencies of Tcf1^+^ cells within lung tumors, and this correlated with their significantly enhanced accumulation within tumors (Fig. 2B). Correspondingly, treatment with cJun.ROR1 CAR-T cells, but not BATF.ROR1 CAR-T cells, induced a modest but significant extension in survival compared to treatment with ROR1 CAR-T cells (Fig. 2C), suggesting that improving preservation of Tcf1^+^ CAR-T cells *in vivo* can improve tumor control.

**Fig. 2.**
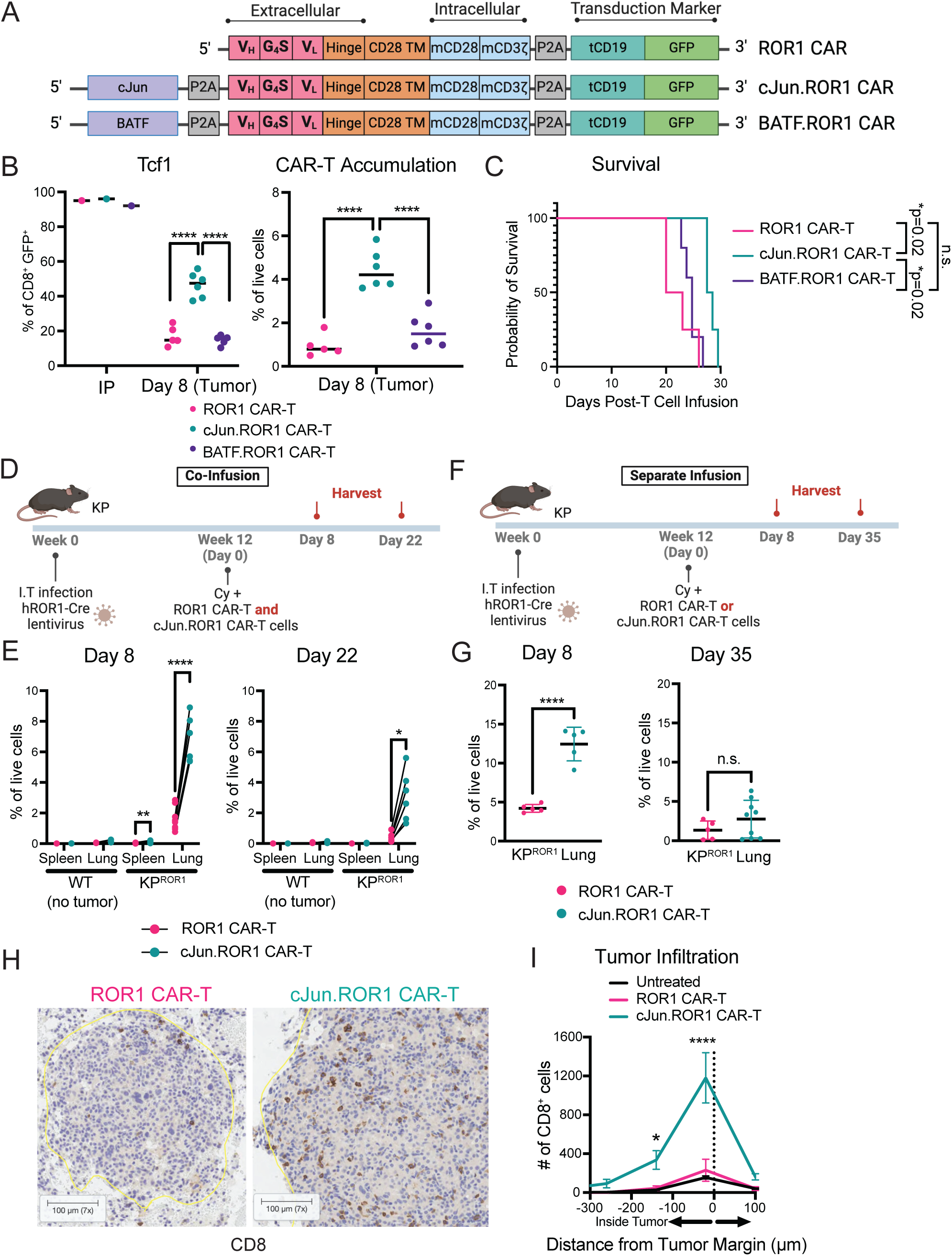
Overexpression of AP-1 family TFs affects maintenance of intratumoral Tcf1^+^ CAR-T cells and long-term persistence *in vivo*. A) Schematic of ROR1 CAR, cJun.ROR1 CAR, and BATF.ROR1 CAR retroviral constructs used to generate murine CAR-T cells. TM = transmembrane. tCD19 = truncated murine CD19. B) Tcf1 expression (left) and accumulation (right) of CD8^+^GFP^+^ CAR-T cells in infusion products (**IP**) and within tumors 8 days post-infusion into B6 mice transplanted with KP-ROR1 lung tumors. One-way ANOVA with Tukey’s post-test. N=5-6 mice per group. C) Survival of B6 mice transplanted with KP-ROR1 lung tumors and treated with Cy and indicated CAR-T cells. Log-rank Mantel-Cox test. N=4-5 mice per groups. D) Treatment scheme showing co-infusion of CAR-T cells. KP = Kras^LSL-G12D/+^p53^f/f^. I.T = intratracheal. Cy = cyclophosphamide. E) Frequency of CD45.1^+^CD8^+^GFP^+^ ROR1 (pink) and CD45.2^+^CD8^+^GFP^+^ cJun.ROR1 CAR-T cells (teal) of total live cells in spleens or lungs from non-tumor bearing WT or tumor-bearing KP^ROR1^ mice 8 and 22 days post-CAR-T cell infusion. Lines indicate paired comparisons between ROR1 CAR-T and cJun.ROR1 CAR-T cells in the same mice. N=3-6 mice per group. Paired Student’s two-way t-test. F) Treatment scheme showing separate CAR-T infusion. G) Frequency of CD8^+^GFP^+^ ROR1 (pink) and cJun.ROR1 CAR-T cells (teal) of total live cells in lungs of tumor-bearing KP^ROR1^ mice 8 and 35 days post-CAR-T cell infusion. N=5-9 mice per group. Unpaired Student’s two-way t-test. H) Representative IHC staining for CD8 on lungs of KP^ROR1^ mice 21 days post-infusion with ROR1 CAR-T or cJun.ROR1 CAR-T cells. Yellow line indicates tumor margin. I) IHC analysis showing quantification of CD8^+^ cells at various distances from tumor margin in KP^ROR1^ lungs 21 days post-infusion of ROR1 CAR-T or cJun.ROR1 CAR-T cells. Two-way ANOVA with Tukey’s post-test. N=3 mice per group. Data are representative of 2-5 independent experiments.

Because Tcf1^+^ Tpex have been shown to sustain and replenish T cells within tumors, we next examined how c-Jun overexpression affected long-term CAR-T cell persistence *in vivo*. Long-term T cell persistence, however, is difficult to assess in transplantable tumor models, in which fully mature tumor cells are implanted and progress rapidly. Thus, we instead evaluated how c-Jun overexpression affected CAR-T cell persistence in the autochthonous KP^ROR1^ model, in which tumors progress from defined oncogenic mutations in single lung epithelial cells over 4-6 months. To directly compare ROR1 CAR-T and cJun.ROR1 CAR-T cells head-to-head, we co-infused a 1:1 ratio of cJun.ROR1 and ROR1 CAR-T cells into tumor-bearing KP^ROR1^ mice (Fig. 2D). By 7 days post-infusion, cJun.ROR1 CAR-T cells significantly outcompeted ROR1 CAR-T cells at accumulating within the same KP^ROR1^ lung tumors (Fig. 2E). This competitive advantage was maintained up to at least 22 days post-infusion, though both cJun.ROR1 and ROR1 CAR-T cells declined in frequency over time. By contrast, both cJun.ROR1 and ROR1 CAR-T cells maintained a ∼1:1 ratio in spleens and lungs of non-tumor-bearing mice and in spleens of KP^ROR1^ mice, suggesting that the accumulation benefit observed with c-Jun overexpression was antigen-dependent. We observed similar trends when cJun.ROR1 and ROR1 CAR-T cells were infused into separate mice, with cJun.ROR1 CAR-T cells showing significantly increased accumulation 8 days post-infusion but no significant difference by 35 days post-infusion (Fig. 2F, 2G), indicating the early benefits of c-Jun overexpression were also evident in a non-competitive setting. Immunohistochemistry of KP^ROR1^ lung tumors showed that cJun.ROR1 CAR-T cells persisted at higher density and penetrated deeper within tumors compared to ROR1 CAR-T cells by 21 days post-infusion (Fig. 2H-2I). Altogether, these data show that c-Jun overexpression significantly enhances the ability of CAR-T cells to establish a PD-1^+^Tcf1^+^ resource population within tumors, and this correlates with improved persistence deeper within tumor nests in a cell-intrinsic and antigen-dependent manner; however, these benefits wane with time and result in only modestly improved survival, indicating the effects of c-Jun overexpression alone are insufficient to substantially enhance CAR-T cell activity in models of lung cancer.

### c-Jun overexpression improves preservation of stem-like PD-1^+^Tcf1^+^ CAR-T cells within lung tumors in a cell-intrinsic manner

Although Tcf1^+^ cells were increased among cJun.ROR1 CAR-T cells within tumors, whether this subset expresses stemness gene signatures comparable to conventional MHC-restricted Tpex is not clear. To determine more comprehensively how c-Jun overexpression affected CAR-T cell differentiation in the TME, we treated tumor-bearing KP^ROR1^ mice with Cy and either cJun.ROR1 or ROR1 CAR-T cells and harvested lung tumors 9 days and 30 days post-infusion. We used single-cell RNA sequencing (**scRNAseq**) to resolve the transcriptional profile of CD8^+^ T cells enriched from tumors at each time point, as well as of cJun.ROR1 or ROR1 CAR-T cells prior to infusion. We recovered some non-T cell populations that likely reflected imperfect purity of the CD8 positive selection protocol (Fig. S3A); additionally, we detected a small subset of cells that exclusively expressed *Cd8, Cd4,* and *Rag1* that likely represented contaminating thymocytes isolated during lung dissection, as this subset was derived from a single mouse (Fig. S3B, S3C). Thus, we re-clustered the data to focus on *Cd3e*-expressing T cells and to exclude contaminating thymocytes (Fig. S3D). The *CAR* transgene was highly expressed in a subset of clusters, which we further sub-clustered to better resolve the transcriptional profile of CAR-T cells (Fig. S3D, S3E).

Unsupervised clustering analysis of CAR-T cells revealed a range of cell states that segregated based on time point (Fig. 3A, 3B). Cells in the CAR-T cell infusion product (“**IP**”) highly expressed stemness-associated genes, such as *Tcf7*, *Il7r*, and *Bach2*, and lacked expression of *Pdcd1* and *Tox*, consistent with a non-exhausted central memory-like (**Tcm**) phenotype prior to infusion (Fig. 3B, 3C). After infusion, a small subset of cells present at both Day 9 and Day 30 retained expression of central memory-like genes similar to the infusion product and likely represented cells that had not yet engaged antigen *in vivo*, which we termed “**Tcm**”. At Day 9, we resolved 3 clusters of cells distinguished by their upregulation of the exhaustion-associated genes *Pdcd1* and *Tox*. The largest of these 3 clusters expressed a combination of memory-associated genes (*Tcf7*, *Il7r*, *Bach2*) and exhaustion-associated genes (*Pdcd1*, *Tox*, *Tigit*, *Lag3*), consistent with the phenotype of PD-1^+^Tcf1^+^ stem-like precursor exhausted cells (**Tpex**) that have been shown to mediate the proliferative burst in response to PD-1/PD-L1 axis blockade (Fig. 3C).^35–42,78^ This “**Tpex**” cluster showed strong enrichment of multiple gene signatures of PD-1^+^Tcf1^+^ Tpex derived from studies in both mouse and human T cells in chronic infection and cancer, suggesting high transcriptional similarity between this CAR-T cell subset and endogenous Tpex cells (Fig. 3D, S3F).^38,53,78,79^ We also identified a cluster of effector-like exhausted cells (“**Teff**”) which expressed inhibitory receptors and markers of exhaustion (*Pdcd1*, *Tox*, *Havcr2*,), lacked expression of memory-like genes, and expressed the highest levels of genes associated with effector function (*Ifng*, *Batf*) and costimulatory and IL-2 signaling (*Tnfsf11*, *Tnfrsf4*, *Cd70*, *Il2ra*, *Stat5a*) (Fig. 3C). The final cluster enriched at Day 9 showed relatively lower expression of memory-associated, effector-like, and costimulatory genes, and higher expression of inhibitory receptor and exhaustion-related genes, suggesting a terminally exhausted-like phenotype (“**Tex D9**”). By Day 30, although a small subset cells were found in the Tcm and Tpex clusters, the majority of T cells were found in a cluster that lacked expression of memory– and costimulatory-related genes and showed the highest expression of inhibitory receptor genes and genes associated with terminal exhaustion (*Tox, Gzmk, Prf1, Eomes, Entpd1*), which we termed “**Tex D30**” (Fig. 3C). This cluster showed strong enrichment for gene signatures of terminal exhaustion, derived from both mouse and human T cells in chronic infection and cancer, that has been associated with failure to respond to PD-L1 blockade (Fig. 3D, Fig. S3G).^38,53,78,79^ Finally, we also resolved a cluster of “**Cycling**” cells that highly expressed the cell cycle-associated genes *Mki67* and *Stmn1* and was comprised of cells from each time point.

**Fig. 3.**
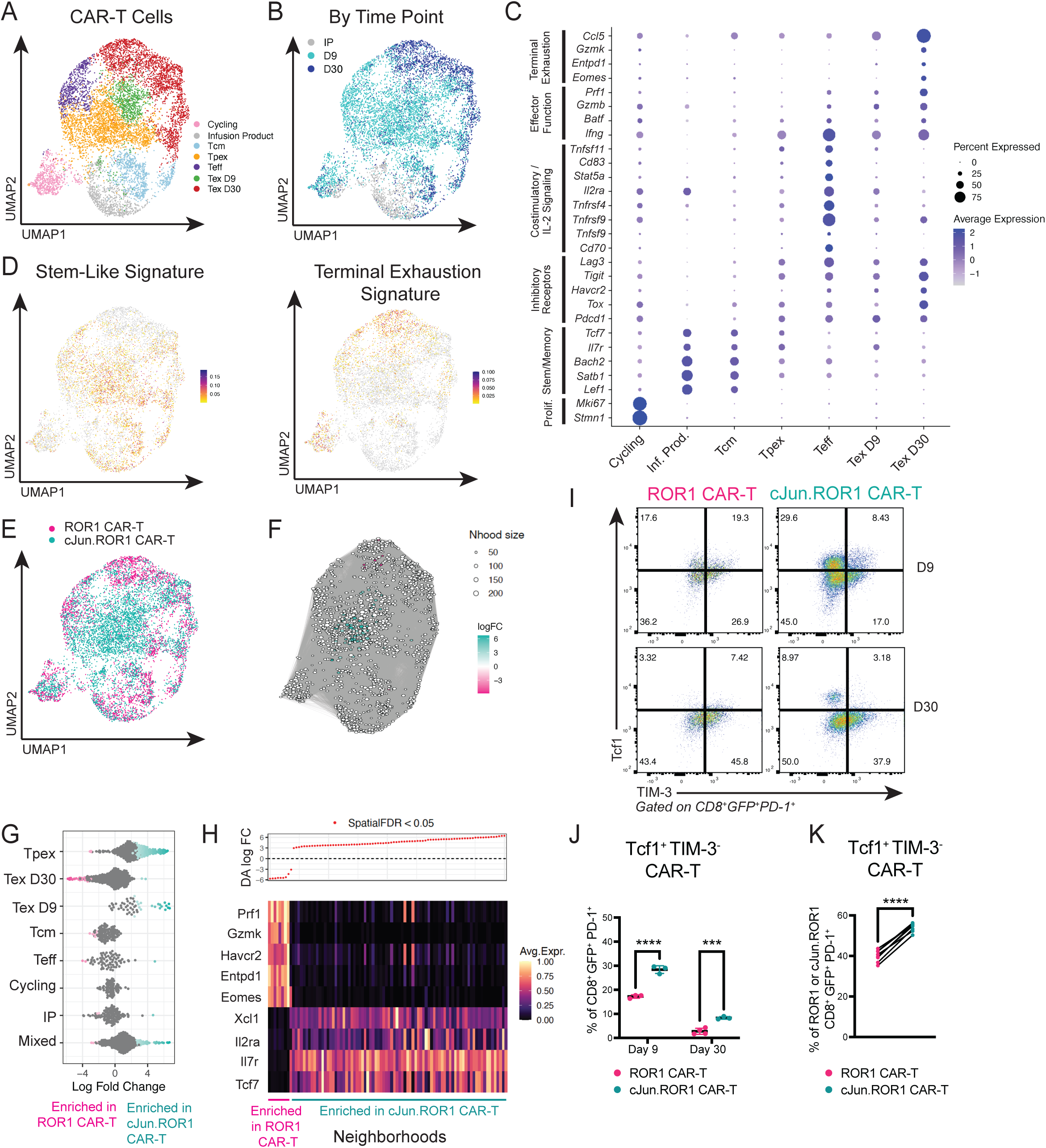
c-Jun overexpression improves preservation of stem-like PD-1^+^Tcf1^+^ CAR-T cells within lung tumors in a cell-intrinsic manner. A) Unsupervised clustering of ROR1 and cJun.ROR1 CAR-T cells from the Infusion Product or from KP^ROR1^ tumors 9 and 30 days post-infusion and analyzed by scRNA-seq. B) Clusters in (A) colored by time point. C) Dot plot showing expression of genes that define each CAR-T cell subcluster. D) Geneset scores of enriched pathways from Jansen et al.^53^ in CAR-T cell subclusters. E) Clusters in (A) colored by CAR-T cell product. F) K-nearest neighbor graph using Milo showing neighborhoods enriched in cJun.ROR1 CAR-Ts (teal) or ROR1 CAR-Ts (pink). G) Differential abundance analysis using Milo showing fold change enrichment of neighborhoods within specific CAR-T subclusters in either cJun.ROR1 CAR-Ts (teal) or ROR1 CAR-Ts (pink). H) Heatmap showing genes differentially expressed between neighborhoods enriched in ROR1 CAR-T or cJun.ROR1 CAR-T cells. I, J) Representative flow cytometry plots (I) and summary (J) showing frequency of Tcf1^+^TIM-3^−^ stem-like subset among CD8^+^GFP^+^PD-1^+^ CAR-T cells in KP^ROR1^ lungs 9 and 30 days post-infusion. N=3-4 mice per group. Two-way ANOVA with Tukey’s post-test. K) Frequency of Tcf1^+^TIM-3^−^stem-like cells among CD45.1^+^CD8^+^GFP^+^PD-1^+^ ROR1 CAR-T (pink) and CD45.2^+^CD8^+^GFP^+^PD-1^+^ cJun.ROR1 CAR-T cells (teal) in KP^ROR1^ lungs 9 days post-co-infusion. Lines indicated matched ROR1 and cJun.ROR1 CAR-T cells within the same mice. N=6 mice per group. Paired Student’s two-way t-test. Data are representative of 2 independent experiments.

To compare the relative abundance of ROR1 CAR-T cells and cJun.ROR1 CAR-T cells across these different cell states, we performed differential abundance testing using Milo,^80^ a method for differential abundance analysis across overlapping “neighborhoods”. Notably, cJun.ROR1 CAR-T cells were significantly enriched among 81 neighborhoods localized within the Tpex cluster and defined by upregulation of the memory-associated genes *Tcf7*, *Il7r*, *Il2ra*, and *Xcl1* (Fig. 3E-H). By contrast, ROR1 CAR-T cells were significantly enriched in 9 neighborhoods localized within the Tex D30 cluster that were defined by upregulation of genes associated with terminal exhaustion (*Eomes, Entpd1, Havcr2, Prf1, Gzmk*), indicating that ROR1 CAR-T cells show more rapid terminal exhaustion while cJun.ROR1 CAR-T cells show improved preservation of stem-like Tpex (Fig. 3E-H).

Consistent with our scRNAseq data, we used flow cytometry to validate that cJun.ROR1 CAR-T cells showed significantly higher frequency of PD-1^+^Tcf1^+^TIM-3^−^ Tpex up to 30 days post-infusion, indicating that c-Jun overexpression enhanced CAR-T differentiation into Tpex *in vivo* (Fig. 3I, 3J). However, Tpex frequency declined over time, suggesting progressive differentiation *in vivo*. Because antigen stimulation can drive progressive differentiation,^45^ we wondered whether the enhanced function of cJun.ROR1 CAR-T cells might drive a reduction in ROR1^+^ tumor burden that could indirectly lead to less stimulation and better preservation of Tpex. To test this, we co-infused cJun.ROR1 and ROR1 CAR-T cells into tumor-bearing KP^ROR1^ mice and compared their phenotype within the same tumors. cJun.ROR1 CAR-T cells maintained a higher proportion of PD-1^+^Tcf1^+^TIM-3^−^ Tpex even in the co-infusion setting, demonstrating that enhanced Tpex preservation is due to cell-intrinsic effects of c-Jun overexpression on CAR-T cells and not due to extrinsic differences in the TME (Fig. 3K). Thus, our data demonstrate that c-Jun overexpression enables CAR-T cells to preserve an intratumoral stem-like subset that transcriptionally resembles endogenous Tpex cells.

### c-Jun.ROR1 CAR-T cells show improved function but ultimately exhaust in vivo

Although c-Jun overexpression enabled CAR-T cells to establish a Tpex reservoir *in vivo*, our scRNAseq data showed that cJun.ROR1 CAR-T cells ultimately upregulated gene signatures associated with exhaustion (including *Pdcd1* and *Tox*), suggesting that c-Jun overexpression alone may be insufficient to prevent exhaustion and that this may limit CAR-T cell activity *in vivo*. To determine how c-Jun overexpression impacted the development of exhaustion in ROR1 CAR-T cells, we analyzed the expression of co-inhibitory receptors on tumor-infiltrating CAR-T cells over time, as higher and sustained levels of PD-1 and other co-inhibitory receptors is associated with T cell exhaustion.^78,81–84^ Nearly all CAR-T cells upregulated and maintained PD-1 expression within KP^ROR1^ tumors, though cJun.ROR1 CAR-T cells upregulated PD-1 to significantly lower levels and showed lower expression of TIM-3 and LAG-3 compared to ROR1 CAR-T cells 9 days post-infusion (Fig. 4A, 4B). Nevertheless, cJun.ROR1 CAR-T cells eventually upregulated PD-1 and TIM-3 to levels comparable to ROR1 CAR-T cells by day 30 post-infusion, with only LAG-3 maintained at significantly lower levels. Likewise, cJun.ROR1 CAR-T cells in KP^ROR1^ tumors showed higher production of IFNγ, TNFα, and IL-2 compared to ROR1 CAR-T cells within 9 days post-infusion. (Fig. 4C, 4D), but this advantage was largely lost by day 35 post-infusion (Fig. 4C, 4D). Compared to ROR1 CAR-Ts, cJun.ROR1 CAR-T cells expressed significantly lower levels of Tox, a transcription factor shown to drive and reinforce commitment to the “exhaustion” program in CD8^+^ T cells (Fig. 4E, 4F)^85–88^; however, cJun.ROR1 CAR-Ts progressively upregulated Tox over time, suggesting the beneficial effects of c-Jun overexpression were incomplete. Notably, c-Jun overexpression showed similarly beneficial, but incomplete, effects in both ROR1.BBζ and ROR1.28ζ CAR-T cells, with all treated mice showing only modestly improved survival and succumbing to disease (Fig. S4). Altogether, these data indicate that c-Jun overexpression is insufficient to prevent the development of exhaustion in ROR1 CAR-T cells *in vivo*, which limits their anti-tumor activity against aggressive lung tumors.

**Fig. 4.**
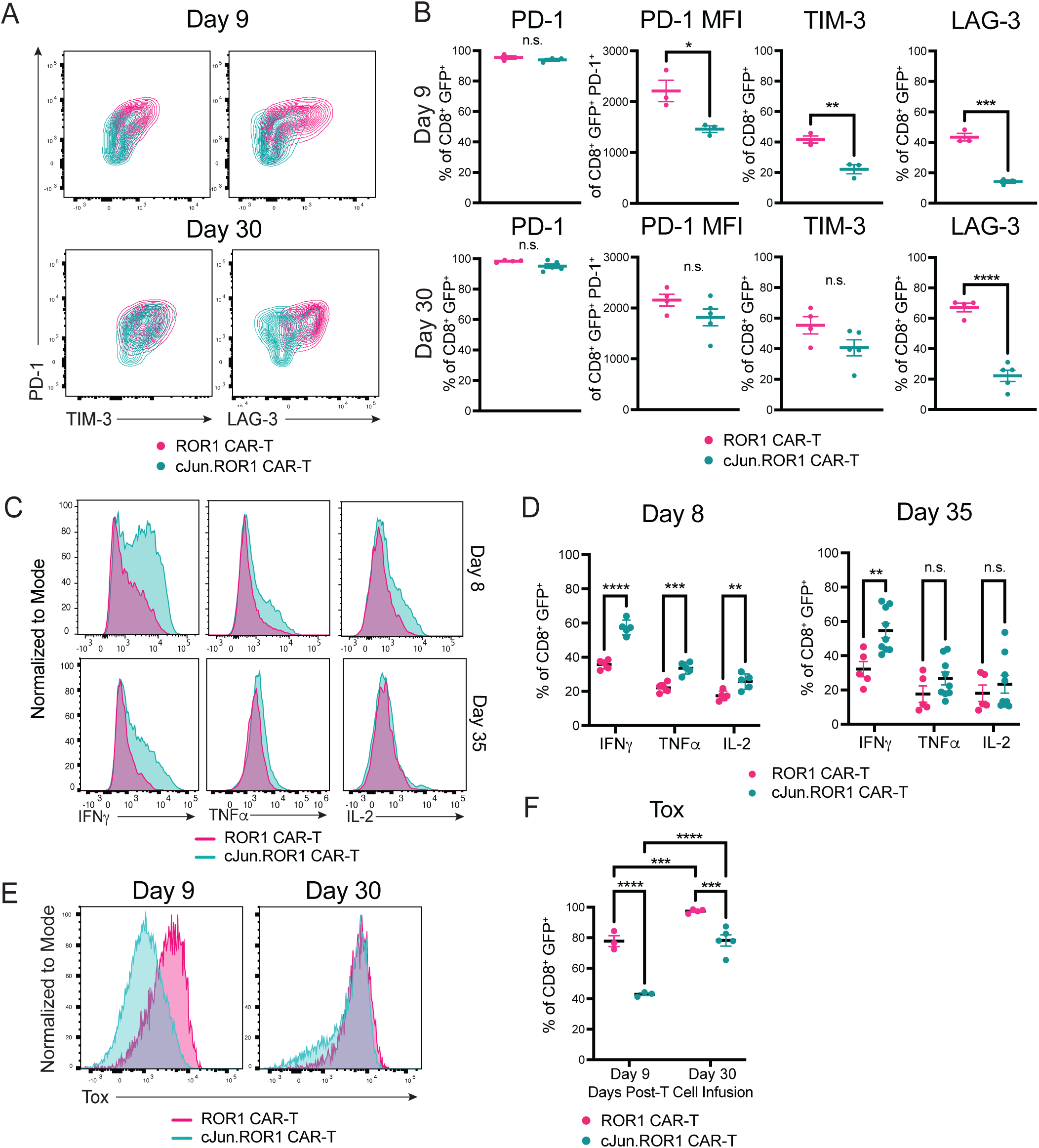
cJun.ROR1 CAR-T cells show improved function but ultimately exhaust *in vivo*. A, B): Representative flow cytometry plots (A) and summary (B) of inhibitory receptor expression on CD8^+^GFP^+^ ROR1 CAR-T (pink) and cJun.ROR1 CAR-T cells (teal) from KP^ROR1^ lungs 9 days and 30 days post-infusion. N=3-5 mice per group. Unpaired Student’s two-way t-test. C, D) Representative flow cytometry plots (C) and summary (D) of intracellular cytokine staining of CD8^+^GFP^+^ ROR1 CAR-T (pink) and cJun.ROR1 CAR-T cells (teal) in KP^ROR1^ lungs 8 and 35 days post-infusion after *ex vivo* restimulation with PMA and ionomycin. N=5-9 mice per group. Unpaired Student’s two-way t-test. E, F) Representative flow cytometry plots (E) and summary (F) of Tox expression in CD8^+^GFP^+^ ROR1 CAR-T (pink) and cJun.ROR1 CAR-T cells (teal) in KP^ROR1^ lungs 9 and 30 days post-infusion. N=3-5 mice per group. Two-way ANOVA with Tukey’s post-test. Data are representative of 2-5 independent experiments.

### c-Jun overexpression dramatically sensitizes ROR1 CAR-T cells to enhancement with PD-L1 blockade

Although c-Jun overexpression alone did not substantially enhance ROR1 CAR-T cell-mediated tumor control, we hypothesized that its ability to enhance numbers of PD-1^+^Tcf1^+^ stem-like CAR-T cells within tumors may sensitize them to enhancement with ICI. To test this, we treated tumor-bearing KP^ROR1^ mice with ROR1 CAR-T or cJun.ROR1 CAR-T cells and administered either blocking anti-PD-L1 antibody or vehicle every 3-4 days (Fig. 5A). By 5 days post-infusion, PD-L1 blockade had no effect on ROR1 CAR-T cell numbers, as observed before, but drastically boosted numbers of tumor-infiltrating cJun.ROR1 CAR-T cells >10-fold. PD-1 expression remained uniformly high in CAR-T cells across all treatment groups but increased slightly in cJun.ROR1 CAR-T cells with PD-L1 blockade, suggesting that this combination drove increased activation of cJun.ROR1 CAR-T cells (Fig.5B, 5C). Interestingly, combination treatment with cJun.ROR1 CAR-T cells and anti-PD-L1 also drove a synergistic increase in numbers of tumor-infiltrating endogenous CD8^+^ T cells, which also showed greater expression of PD-1, suggesting that treatment with c-Jun.ROR1 CAR-T cells may also sensitize endogenous CD8^+^ T cells to enhancement with PD-L1 blockade (Fig. 5C). Whereas PD-L1 blockade drove terminal differentiation of ROR1 CAR-T cells (Fig 5D, 5E), it increased frequencies of both PD-1^+^Tcf1^+^TIM-3^−^ Tpex and more differentiated PD-1^+^Tcf1^−^TIM-3^+^ Tex cells among cJun.ROR1 CAR-T cells, suggesting an ability to both expand a stem-like reservoir and generate more differentiated effector cells. Thus, c-Jun overexpression enhances ROR1 CAR-T accumulation and protects them from terminal differentiation in response to PD-L1 blockade.

**Fig. 5.**
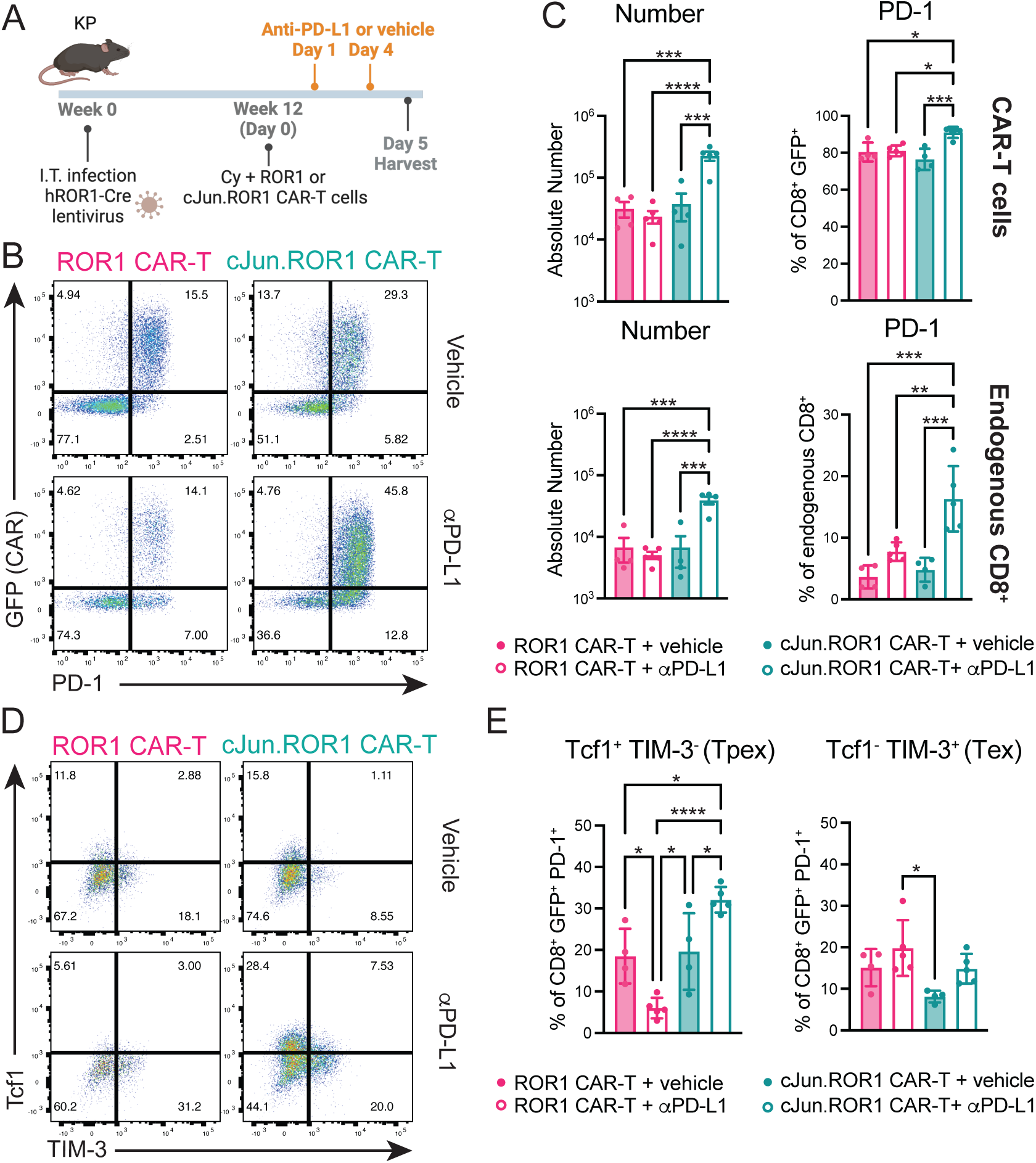
c-Jun overexpression dramatically sensitizes ROR1 CAR-T cells to enhancement with PD-L1 blockade. A) Treatment schematic. I.T. = intratracheal. Cy = cyclophosphamide. B) Representative flow cytometry plots showing expression of the CAR transduction marker (GFP) and PD-1 in CD8^+^ T cells from KP^ROR1^ lungs 5 days post-infusion. C) Absolute number and PD-1 expression in CD8^+^GFP^+^ CAR-T cells (top) and CD8^+^GFP^−^ endogenous T cells (bottom) in lungs of KP^ROR1^ mice treated as indicated. N=4-5 mice per group. One-way ANOVA with Tukey’s post-test. D) Representative flow cytometry plots showing Tcf1 and TIM-3 expression in CD8^+^GFP^+^PD-1^+^ CAR-T cells in KP^ROR1^ lungs 5 days post-infusion. E) Frequency of Tcf1^+^TIM-3^−^ stem-like precursor exhausted (**Tpex**) and Tcf1^−^TIM-3^+^ terminally exhausted (**Tex**) subsets of CD8^+^GFP^+^PD-1^+^ CAR-T cells in KP^ROR1^ lungs 5 days post-infusion. N=4-5 mice per group. One-way ANOVA with Tukey’s post-test. Data are representative of 2 independent experiments

### c-Jun levels are downregulated post-transcriptionally in cJun.ROR1 CAR-T cells and restored by PD-L1 blockade

c-Jun overexpression was previously shown to prevent CAR-T cell exhaustion by competing with and displacing inhibitory AP-1 transcription factors from chromatin, suggesting that c-Jun levels must be sufficiently high relative to inhibitory transcription factors to mediate successful reprogramming of CAR-T cells.^65^ We hypothesized that incomplete prevention of exhaustion in cJun.ROR1 CAR-T cells may be due to insufficiently high overexpression of c-Jun. To test this, we used an antibody that detects both endogenous and transgenic murine c-Jun and used flow cytometry to measure c-Jun expression in CAR-T cells over time. Whereas cJun.ROR1 CAR-T cells uniformly expressed higher levels of c-Jun relative to ROR1 CAR-T cells prior to infusion, they significantly downregulated c-Jun levels in KP^ROR1^ tumors as early as 9 days post-infusion (Fig. 6A, 6B).

**Fig. 6.**
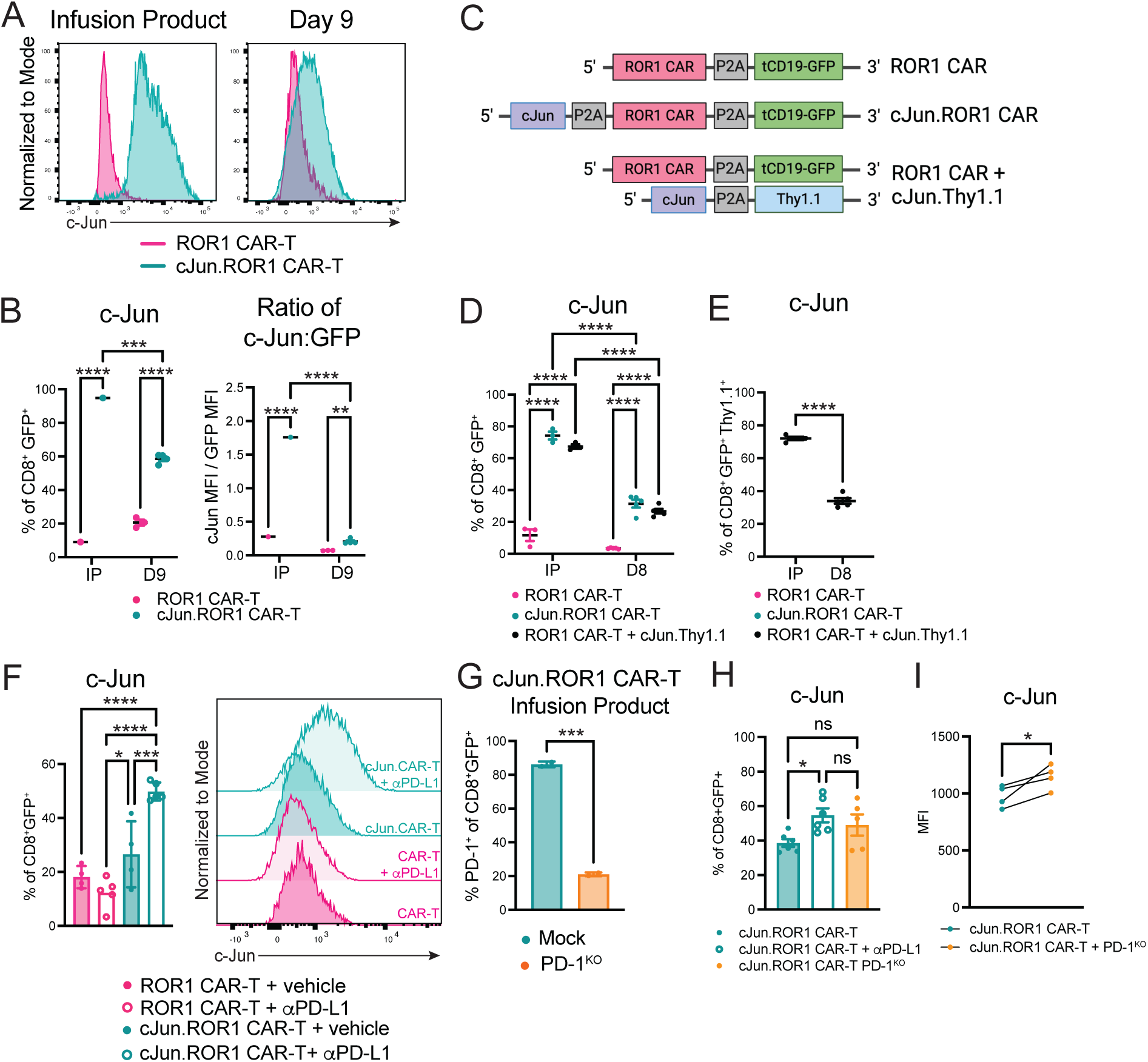
c-Jun levels are downregulated post-transcriptionally in cJun.ROR1 CAR-T cells and restored by PD-L1 blockade. A) Flow cytometry plots showing murine c-Jun expression in ROR1 CAR-T (pink) and cJun.ROR1 CAR-T cells (teal) pre-infusion and within KP^ROR1^ lungs 9 days post-infusion. B) Frequency of c-Jun (left) and ratio of c-Jun to GFP median fluorescence intensity (**MFI**) (right) in CD8^+^GFP^+^ CAR-T cells in the infusion product (**IP**) and in KP^ROR1^ lungs 9 days post-infusion. N=3-5 mice per group. Two-way ANOVA with Tukey’s post-test. C) Schematic of retroviral CAR and c-Jun constructs. D) Total c-Jun expression in CD8^+^GFP^+^ CAR-T cells in the IP or in KP^ROR1^ lungs 8 days post-infusion. N=5 mice per group. Two-way ANOVA with Tukey’s post-test. E) Total c-Jun expression in CD8^+^GFP^+^Thy1.1^+^ CAR-T cells co-transduced with ROR1 CAR-P2A-tCD19/GFP and cJun-P2A-Thy1.1 vectors in the IP and in KP^ROR1^ lungs 8 days post-infusion. N=5 mice per group. Unpaired Student’s two-way t-test. F) Summary (left) and representative flow cytometry plot (right) of c-Jun expression in CD8^+^GFP^+^ ROR1 CAR-T (pink) and cJun.ROR1 CAR-T cells (teal) in KP^ROR1^ lungs 5 days post-infusion and treatment with vehicle (filled) or αPD-L1 antibody (unfilled). N=4-5 mice per group. One-way ANOVA with Tukey’s post-test. G) PD-1 expression on CRISPR/Cas9 mock-edited and PD-1^KO^ cJun.ROR1 CAR-T cell infusion products 24 hr post-stimulation with PMA/ionomycin. N=3 technical replicates. Unpaired Student’s two-way t-test. H) c-Jun levels in tumor-infiltrating cJun.ROR1 CAR-T cells treated as indicated 9 days post-infusion into B6 mice bearing transplanted KP^ROR1^ tumors. N=5-6 mice per group. One-way ANOVA with Tukey’s post-test. I) c-Jun levels in mock-edited and PD-1^KO^ cJun.ROR1 CAR-T cells 9 days post-co-infusion into tumor-bearing KP^ROR1^ mice. N=4 mice per group. Paired Student’s two-way t-test. Data are representative of 2 independent experiments.

Downregulation of c-Jun could be due to decreased transcription and/or epigenetic silencing of the cJun-P2A-CAR-P2A-tCD19/GFP transgene, as retroviral vectors have been reported to undergo DNA methylation in T cells *in vivo*.^89^ We hypothesized that this would result in equivalent downregulation of both c-Jun and the inert tCD19/GFP transduction marker encoded by the retroviral transgene, such that the ratio of c-Jun to GFP protein would remain equivalent. Instead, we found that the ratio of c-Jun to GFP levels decreased significantly in cJun.ROR1 CAR-T cells within KP^ROR1^ tumors 9 days post-infusion relative to pre-infusion, suggesting that c-Jun was being selectively downregulated relative to the tCD19/GFP marker (Fig. 6B).

Alternatively, high CAR expression has been shown to drive activation-induced cell death^90^, and because c-Jun is co-expressed with the ROR1 CAR, death of T cells with high CAR (and thus high c-Jun) expression may leave behind a residual CAR-T population with lower c-Jun levels. To determine whether c-Jun was selectively downregulated independently of the ROR1 CAR, we designed constructs which unlinked expression of c-Jun and the ROR1 CAR. We co-transduced T cells with CAR-P2A-tCD19/GFP and cJun-P2A-Thy1.1 vectors and compared them to T cells transduced with our standard cJun-P2A-CAR-P2A-tCD19/GFP vector (Fig. 6C). Both CAR-T cells transduced with a single cJun.ROR1 CAR vector or co-transduced with separate c-Jun and ROR1 CAR vectors showed equivalently high c-Jun expression prior to infusion and similar downregulation of c-Jun in KP^ROR1^ tumors within 8 days post-infusion, indicating that c-Jun downregulation occurred independently of co-expression with the ROR1 CAR (Fig. 6D). Interestingly, although c-Jun and the Thy.1 transduction marker were highly co-expressed in the pre-infusion product, <40% of Thy1.1^+^ CAR-T cells maintained c-Jun expression in KP^ROR1^ tumors 8 days post-infusion (Fig. 6E). Thus, tumor-infiltrating cJun.ROR1 CAR-T cells downregulate c-Jun independently of the ROR1 CAR and Thy1.1 or tCD19/GFP markers, suggesting that c-Jun downregulation occurs post-transcriptionally.

c-Jun stability is regulated in part by phosphorylation by MAPK family kinases,^91^ and TCR signaling induces phosphorylation of c-Jun at sites that protect it from ubiquitin-mediated degradation and increase its half-life.^92–96^ Because of its ability to inhibit proximal TCR signaling, PD-1 signaling can inhibit c-Jun phosphorylation and restrain AP-1 activity in T cell lymphomas.^97^ Given that PD-L1 blockade dramatically increased c-Jun CAR-T cells within KP^ROR1^ tumors, we wondered whether PD-1/PD-L1 signaling was responsible for reduced c-Jun levels in cJun.ROR1 CAR-T cells. We treated tumor-bearing KP^ROR1^ mice with cJun.ROR1 or ROR1 CAR-T cells and administered anti-PD-L1 or vehicle as before. By 5 days post-infusion, c-Jun levels were only slightly, but insignificantly, elevated in cJun.ROR1 CAR-T cells relative to ROR1 CAR-T cells (Fig. 6F). However, treatment with anti-PD-L1 dramatically increased c-Jun levels in cJun.ROR1 CAR-T cells but had no effect on ROR1 CAR-T cells (Fig. 6F), indicating that c-Jun downregulation was PD-L1-dependent.

The effects of PD-L1 blockade on c-Jun levels in CAR-T cells could be due to direct effects of disrupting PD-1 signaling in CAR-T cells and/or due to indirect effects on other PD-1^+^ or PD-L1^+^ cells in the TME.^77–80^ To determine whether intrinsic PD-1 signaling in CAR-T cells was responsible for reduced c-Jun levels, we used CRISPR/Cas9 to selectively knockout PD-1 in CD8^+^ cJun.ROR1 CAR-T cells (**cJun.ROR1^PD^**^1^**^−KO^**) (Fig. 6G). While anti-PD-L1 drove a significant increase in c-Jun levels in c-Jun.ROR1 CAR-T cells, cJun.ROR1^PD1−KO^ CAR-T cells showed intermediate c-Jun levels that were slightly, but insignificantly, increased compared to mock-edited cJun.ROR1 CAR-T cells (Fig. 6H). To control for extrinsic differences in the TME that could influence c-Jun levels, we compared c-Jun levels in mock-edited and PD-1^KO^ cJun.ROR1 CAR-T cells co-infused into the same tumor-bearing KP^ROR1^ mice. Within the same tumors, PD-1^KO^ cJun.ROR1 CAR-T cells showed significantly increased c-Jun levels relative to mock-edited cells, though the magnitude of increase was not as large as that observed with PD-L1 blockade (Fig. 6I). These data suggest that c-Jun downregulation partially depends on intrinsic PD-1 signaling in cJun.ROR1 CAR-T cells, but that extrinsic effects of anti-PD-L1 on other cell types likely also contribute to c-Jun downregulation.

### cJun.ROR1 CAR-T cells localize more closely to PD-L1^+^ cells and show reduced terminal differentiation in response to PD-L1 blockade

One possible explanation for the enhanced sensitivity of cJun.ROR1 CAR-T cells to PD-L1 blockade could be increased PD-1/PD-L1 signaling in tumors treated with cJun.ROR1 CAR-T cells. Using scRNAseq data from live cells sorted from CAR-T cell-treated KP^ROR1^ tumors, we found that among all immune cells identified, *Cd274* (which encodes PD-L1) was upregulated to the greatest degree on tumor macrophages and neutrophils, which comprise the dominant immune infiltrate in KP tumors (Fig. S5A-F).^98^ Using flow cytometry, we confirmed that treatment with ROR1 CAR-T cells and cJun.ROR1 CAR-T cells drove similarly high upregulation of PD-L1 on tumor cells, macrophages, and neutrophils by 9 days post-infusion compared to untreated tumors (Fig. S5G), suggesting that differences in PD-L1 upregulation did not explain the vast difference in CAR-T response to PD-L1 blockade.

To resolve whether ROR1 CAR-T and cJun.ROR1 CAR-T cells showed differential localization relative to various endogenous PD-L1^+^ cell types, we used the 10X Xenium spatial transcriptomics platform and designed a custom panel that included probes for genes defining various endogenous immune and stromal cells identified in our scRNA-seq data from KP^ROR1^ tumors (Fig. 3C, S5D). We also included probes for the ROR1 CAR transgene and ROR1-Cre lentiviral vector, enabling us to specifically identify *CAR^+^* T cells and *ROR1*^+^ tumor cells. We treated tumor-bearing KP^ROR1^ mice with ROR1 CAR-T or cJun.ROR1 CAR-T cells, administering either vehicle or anti-PD-L1 as before, and harvested lungs from each group 7 days post-treatment to analyze by 10X Xenium. Unsupervised clustering enabled us to identify most of the cell types previously identified by scRNA-seq, including subsets of T cells, B cells, macrophages, and neutrophils (Fig. 7A, Fig. S6).

**Fig. 7.**
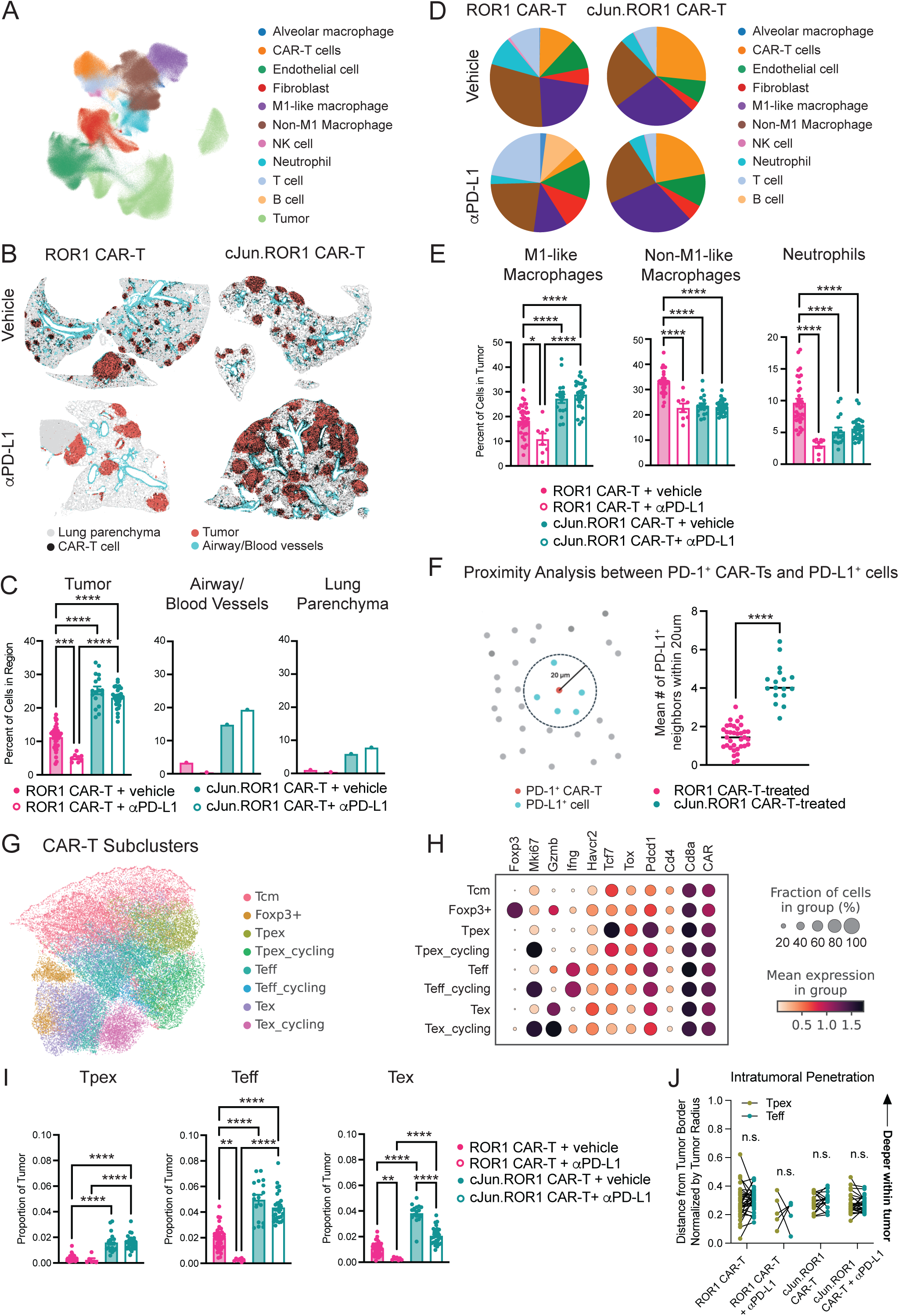
cJun.ROR1 CAR-Ts localize more closely to PD-L1^+^ cells and show reduced terminal differentiation in response to PD-L1 blockade. A) Unsupervised clustering of spatial transcriptomic profiles of cells from lungs of KP^ROR1^ mice 7 days post-treatment with ROR1 CAR-T or cJun.ROR1 CAR-T cells +/− anti-PD-L1. N=1 lung per treatment group. B) Spatial plot showing localization of *ROR1_CART*^+^ cells (black) in tumor (red), airway/blood vessels (blue), and normal lung parenchyma (grey). C) Proportion of cells in indicated spatial regions of each sample expressing the *ROR1_CART* transgene. CAR-T frequency was calculated within each individual tumor nodule. N=8-35 tumor nodules from 1 lung per group. One-way ANOVA with Tukey’s post-test. D) Pie chart showing proportion of each cluster in (A) within tumor region of each sample. E) Left: Proportion of various myeloid clusters among total cells in each tumor nodule. N=8-35 tumor nodules from 1 lung per group. One-way ANOVA with Tukey’s post-test. F) Left: Representative image illustrating proximity analysis calculation. Red = cells co-expressing *ROR1_CART* and *Pdcd1* (PD-1^+^ CAR-T cells). Blue = cells expressing *Cd274* (PD-L1^+^ cells). Right: mean number of PD-L1^+^ neighbors within a 20um radius of a PD-1^+^ CAR-T cell in each tumor nodule. N=17-35 tumor nodules from 1 lung per group. G) Sub-clustering of CAR-T cells. H) Dot plot showing expression of genes defining each cluster. I) Proportion of indicated CAR-T subcluster among total cell within each tumor nodule. N=8-35 tumor nodules from 1 lung per group. One-way ANOVA with Tukey’s post-test. J) Distance of indicated CAR-T subset from tumor border normalized by tumor radius. Connecting lines indicate Tpex and Teff CAR-T subsets within the same tumor nodule. Paired Student’s two-way t-test. N=8-35 tumor nodules from 1 lung per group.

Using both the gene expression profiles and spatial coordinates of each cell, we used Leiden clustering to classify each lung into three distinct spatial regions: normal lung parenchyma, tumor, and a composite non-tumor region that included airway epithelia and blood vessels near tumor nodules (Fig. 7B). In vehicle-treated mice, ROR1 CAR-T cells were enriched within tumors, comprising ∼10% of total cells within this region compared to <3% within non-tumor regions (Fig. 7B, 7C). cJun.ROR1 CAR-T cells were increased in frequency across all three spatial regions but showed preferential enrichment within tumors. Interestingly, anti-PD-L1 treatment induced a dramatic decline in ROR1 CAR-T cells across all three spatial regions, consistent with our prior data (Fig. 1D). By contrast, cJun.ROR1 CAR-T cells maintained elevated levels within tumors upon anti-PD-L1 treatment and instead showed an increase in frequency within the composite airway/blood vessel region, but not the normal lung parenchyma, that may be suggestive of increased extravasation from the circulation (Fig. 7B, 7C).

To understand how ROR1 CAR-T cells and cJun.ROR1 CAR-T cells might differ in their interactions with PD-L1^+^ endogenous cells in the TME, we mapped the localization of each cell type identified via clustering. In all treated mice, the dominant immune infiltrate within tumors was comprised of three subsets of myeloid cells that highly expressed *Cd274* (PD-L1) (Fig. 7D, S6). We identified these myeloid clusters as: 1) “M1-like macrophages”, with higher *Nos2* and lower *Tgfb1* expression; 2) “non-M1-like macrophages”, with higher *Tgfb1* and lower *Nos2* expression; and 3) neutrophils, defined by high expression of *Cxcr2, C5ar1*, and *Mmp9* (Fig. 7D, S6). Neutrophils and non-M1-like macrophages were present at highest frequency in tumors treated with ROR1 CAR-T cells; however, treatment with cJun.ROR1 CAR-T cells induced a decrease in neutrophils and non-M1-like macrophages and an increase in M1-like macrophages within tumors (Fig. 7E). As M1-like macrophages have been reported to exert anti-tumor functions,^99–103^ these changes suggest a polarization of the TME towards a more pro-inflammatory state. Whereas addition of anti-PD-L1 to ROR1 CAR-T therapy induced a decline in M1-like macrophages, addition of anti-PD-L1 to cJun.ROR1 CAR-T therapy maintained elevated frequencies of M1-like macrophages, indicating that cJun.ROR1 CAR-T cells were required to enable the favorable TME-remodeling effects of anti-PD-L1 on tumor macrophages.

We next asked whether the enhanced response of cJun.ROR1 CAR-T cells to PD-L1 blockade might be driven in part by increased interactions of PD-1^+^ CAR-T cells with PD-L1^+^ cells within tumors. Although *Cd274*-expressing macrophages and neutrophils made up a similar fraction (∼60%) of all cells within tumors of mice treated with either ROR1 CAR-T cells or cJun.ROR1 CAR-T cells (Fig. 7D, S6), proximity analysis revealed that PD-1^+^ cJun.CAR-T cells were surrounded by significantly more PD-L1^+^ cells within a 20 um radius than PD-1^+^ ROR1 CAR-T cells (Fig. 7F). These data suggest that altered spatial distribution of PD-L1^+^ cells in closer proximity to cJun.ROR1 CAR-T cells may contribute to their enhanced sensitivity to enhancement with PD-L1 blockade.

Finally, endogenous Tpex cells have been shown to be preferentially enriched in tumor-draining lymph nodes and perivascular APC niches, while more terminally differentiated exhausted T cells localize deeper within tumor nests.^51,53,57^ Because CAR-T cells are not expected to be activated by MHC-dependent interactions with APCs, we wondered whether the Tpex subset formed by cJun.ROR1 or ROR1 CAR-T cells showed distinct localization from more terminally differentiated subsets or whether they were able to persist deep within tumors. To compare localization of Tpex CAR-T cells to other CAR-T subsets, we sub-clustered CAR-T cells based on expression of genes that were most differentially expressed between CAR-T sub-clusters identified from our scRNA-seq dataset (Fig. 3A, 3C). We identified subclusters that correlated closely in gene expression with Tcm (*Pdcd1*^low^ *Tcf7*^hi^), Tpex (*Pdcd1*^hi^ *Tcf7*^hi^ *Havcr2*^lo^), Teff (*Pdcd1*^hi^ *Tcf7*^lo^ *Havcr2*^hi^ *Ifng*^hi^), and Tex (*Pdcd1*^hi^ *Tcf7*^lo^ *Havcr2*^hi^ *Ifng*^lo^ *Gzmb*^hi^) subsets identified by scRNA-seq (Fig. 7G, 7H). Treatment with cJun.ROR1 CAR-T cells induced an increase in the frequency of Tpex, as well as Teff and Tex, subsets within tumors compared to treatment with ROR1 CAR-T cells (Fig. 7I), consistent with our flow and scRNA-seq data. Whereas anti-PD-L1 drove a similar decline in all ROR1 CAR-T cell subsets, it preferentially maintained Tpex and Teff subsets and drove a decline in the Tex subset among cJun.ROR1 CAR-Ts, suggesting enhanced function of cJun.ROR1 CAR-Ts. Interestingly, Tpex cells among both ROR1 CAR-T cells and cJun.ROR1 CAR-T cells showed comparable intratumoral penetration to more differentiated Teff cells within the same tumors, suggesting that CAR-T Tpex are not spatially segregated from Teff and are capable of persisting deep within tumors. Altogether, these data suggest that c-Jun overexpression protects ROR1 CAR-T cells from terminal exhaustion and attrition induced by PD-L1 blockade.

### c-Jun overexpression overcomes resistance of ROR1 CAR-T cells to PD-L1 blockade

To determine whether combination therapy with cJun.ROR1 CAR-T cells and PD-L1 blockade was sufficient to enhance tumor control, we treated tumor-bearing KP^ROR1^ mice with ROR1 CAR-T or cJun.ROR1 CAR-T cells and administered either vehicle or anti-PD-L1 every 3-4 days for a total of 14 days (5 total doses) (Fig. 8A). Four weeks after cessation of anti-PD-L1 treatment, we harvested lungs and used immunohistochemistry to quantify ROR1^+^ tumor burden. Treatment with ROR1 CAR-T cells showed minimal reduction in ROR1^+^ tumor burden relative to untreated KP^ROR1^ mice, with the majority of tumors maintaining ROR1 expression (Fig. 8B, 8C). Treatment with cJun.ROR1 CAR-T cells alone showed a slight but insignificant decrease in ROR1^+^ tumor burden compared to treatment with ROR1 CAR-T cells, consistent with the transient improvement in cJun.ROR1 CAR-T cell function and the modest improvement in survival we observed in transplantable models of lung cancer. Whereas PD-L1 blockade had no significant effect on tumor control by ROR1 CAR-T cells and even slightly worsened it (Fig. 8C), it dramatically improved tumor control by cJun.ROR1 CAR-T cells, with ROR1^+^ tumor eliminated in nearly all treated mice. Total tumor was also significantly decreased in mice treated with cJun.ROR1 CAR-T cells and anti-PD-L1 relative to mice treated with ROR1 CAR-T cells and anti-PD-L1 (Fig. 8C). However, ROR1^−^ tumor continued to progress in mice treated with cJun.ROR1 CAR-T cells and anti-PD-L1, suggesting escape of antigen-null tumor may limit efficacy.

**Fig. 8.**
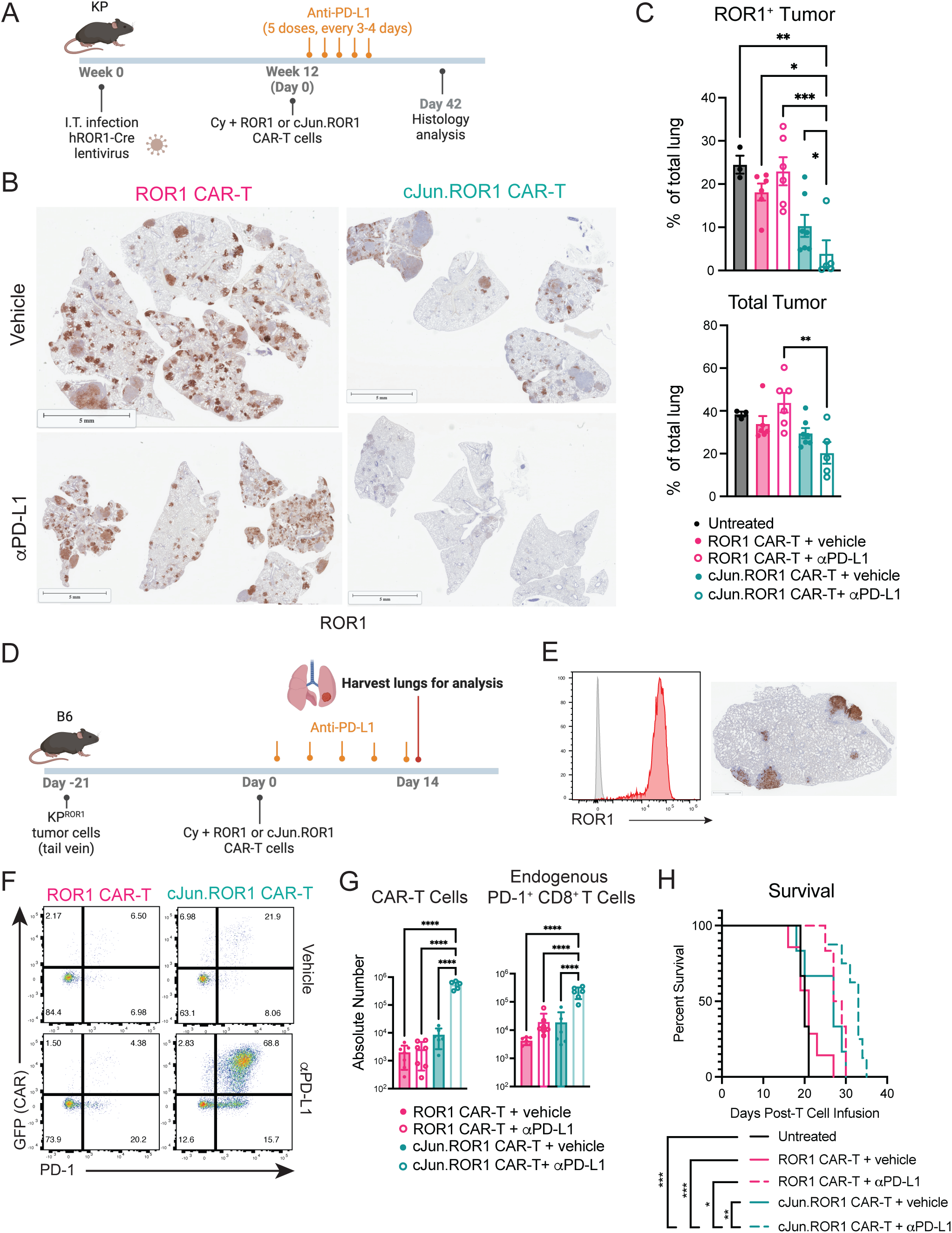
c-Jun overexpression overcomes resistance of ROR1 CAR-T cells to PD-L1 blockade. A) Treatment schematic. I.T. = intratracheal. Cy = cyclophosphamide. B) Representative IHC staining for ROR1 on lungs of KP^ROR1^ mice 42 days post-infusion with ROR1 CAR-T or cJun.ROR1 CAR-T cells. +/− anti-PD-L1. C) Quantification of ROR1^+^ tumor (top) and total tumor (bottom) from IHC images of KP^ROR1^ lungs treated as indicated 42 days post-infusion. N=3-7 mice per group. One-way with ANOVA with Tukey’s post-test. D) Treatment schematic. E) Left: ROR1 expression on KP-ROR1 tumor cell lines derived from GEM model and sorted to be >95% ROR1^+^. Right: representative IHC showing ROR1 expression on lungs of B6 mice transplanted with KP-ROR1 lung tumors 21 days post-tumor injection and pre-CAR-T cell infusion. F) Representative flow plots showing GFP and PD-1 expression on CD8^+^ T cells within lung tumors 14 days post-infusion. G) Absolute number of CD8^+^GFP^+^ CAR-T cells and CD8^+^PD-1^+^ endogenous T cells within lung tumors 14 days post-infusion. N=6-7 mice per group. One-way ANOVA with Tukey’s post-test. H) Survival. Log-rank Mantel-Cox test. N=7-8 mice per group. Data are representative of 2 independent experiments.

To test whether escape of ROR1^null^ tumor is the major factor limiting efficacy of cJun.ROR1 CAR-T and anti-PD-L1 combination therapy, we evaluated this treatment regimen in a lung tumor model with less pre-existing heterogeneity and more uniform expression of ROR1. We induced lung tumor development in B6 mice by intravenously injecting a KP-ROR1 cell lined derived from the GEM model which had been sorted to ensure >95% positivity for ROR1 (Fig. 8D, 8E). Strikingly, in this setting with more uniform ROR1 expression on tumors, PD-L1 blockade again had no effect on numbers of tumor-infiltrating ROR1 CAR-T cells but boosted cJun.ROR1 CAR-T cells numbers within tumors ∼100-fold (Fig. 8F, 8G). Although endogenous CD8^+^PD-1^+^ T cells were increased ∼5-fold by treatment with ROR1 CAR-T cells and anti-PD-L1 together compared to treatment with ROR1 CAR-T cells alone, their numbers increased ∼100 fold upon treatment with cJun.ROR1 CAR-T cells and anti-PD-L1, suggesting a synergistic increase in activation of endogenous T cells by this combination. In contrast to results in the GEM model, addition of anti-PD-L1 to ROR1 CAR-T cell treatment showed a slight improvement in survival, though these effects were likely driven by increases in endogenous CD8^+^ T cells, as numbers of ROR1 CAR-T cells were unchanged by anti-PD-L1 treatment (Fig. 8F, 8G). However, treatment with cJun.ROR1 CAR-T cells and anti-PD-L1 together significantly improved survival compared to all other treatment groups (Fig. 8H). Altogether, our data demonstrate that c-Jun overexpression dramatically reverses CAR-T cell resistance to PD-L1 blockade, enabling efficient regression of ROR1^+^ tumors and improved survival in aggressive models of ROR1^+^ lung cancer.

## DISCUSSION

PD-1^+^Tcf1^+^ Tpex cells are critical for sustaining endogenous anti-tumor immune responses to PD-1/PD-L1 blockade, but whether CAR-T cells can maintain this critical resource population *in vivo* and how this affects their sensitivity to ICI have not been well-defined. Here, we demonstrate that, despite showing high Tcf1 expression, polyfunctionality, and expression of stemness gene signatures prior to infusion, ROR1 CAR-T cells rapidly downregulate Tcf1 and are unable to establish a PD-1^+^Tcf1^+^ stem-like reservoir *in vivo*, resulting in rapid terminal exhaustion and faster attrition in response to PD-L1 blockade. Overexpression of the AP-1 family TF c-Jun, but not BATF, significantly improved preservation of PD-1^+^Tcf1^+^ stem-like CAR-T cells within tumors in a cell-intrinsic manner and sensitized CAR-T cells to enhancement with ICI, enabling nearly complete elimination of ROR1^+^ tumor and improved survival in highly aggressive models of ROR1^+^ lung cancer. Notably, c-Jun overexpression alone was insufficient to prevent CAR-T exhaustion in the lung TME, in contrast to prior work in xenograft models, with progressive CAR-T dysfunction correlated with PD-1-dependent downregulation of c-Jun. PD-L1 blockade restored c-Jun levels, suggesting that PD-1 axis inhibition may be necessary to reveal the full potential of c-Jun-overexpressing CAR-T cells in the suppressive microenvironment of aggressive solid tumors like lung cancer. Altogether, our data show that ROR1 CAR-Ts fail to establish an intratumoral PD-1^+^Tcf1^+^ stem-like reservoir, but that c-Jun overexpression can overcome this defect and may serve as a readily translatable strategy to sensitize previously refractory CAR-Ts to PD-1 axis blockade.

Although ICI can rescue endogenous anti-tumor activity and has been transformative for the treatment of many solid tumors,^104,105^ combination of CAR-T therapy with ICI has shown mixed result in patients, and the mechanisms underlying CAR-T resistance to ICI are unclear.^24–34^ Our data suggest that the ability of CAR-T cells to establish and maintain a PD-1^+^Tcf1^+^ stem-like reservoir *in vivo* is a critical determinant of sensitivity to ICI. Although Tcf1 expression and stemness in cell therapy infusion products is associated with improved clinical responses,^71–77^ signals within the TME may promote terminal differentiation of infused cells and fail to protect a subset of stem-like CAR-T cells that can sustain anti-tumor responses and expand in response to PD-1/PD-L1 blockade. Indeed, signals known to protect endogenous stem-like Tpex cells, such as MHC-I-dependent interactions with DCs in APC niches and tumor-draining lymph nodes,^51,52,57^ are likely bypassed by CAR-T cells, which do not depend on peptide/MHC for activation. Thus, CAR-T cells may be uniquely disadvantaged at establishing this critical stem-like reservoir *in vivo*, which may contribute to their poor persistence, exhaustion, and resistance to PD-L1 blockade.^47^ Indeed, despite showing high Tcf1 expression and polyfunctionality prior to infusion, ROR1 CAR-T cells rapidly downregulated Tcf1 *in vivo* and showed increased terminal differentiation and more rapid attrition in response to PD-L1 blockade, potentially even worsening tumor control. These results mirror those shown in recent studies where depletion of DCs or knockout of MHC-I expression on DCs drove terminal differentiation of Tpex cells and loss of viral control,^51,57^ suggesting that absence of MHC-I-dependent interactions with DCs may drive rapid terminal differentiation of CAR-T cells *in vivo*. Importantly, our data suggest that overexpression of c-Jun may be one strategy that bypasses these limitations and enables CAR-T cells to establish an intratumoral stem-like reservoir that can sensitize them to enhancement with ICI. Notably, c-Jun overexpression did not substantially increase Tcf1 expression or functionality of ROR1 CAR-T cells prior to infusion; however, it may protect CAR-T cells from signals they receive in the TME that promote their terminal differentiation and downregulation of Tcf1. Interestingly, cJun.ROR1 CAR-T cells showed significantly better preservation of Tpex within tumors even when co-infused with ROR1 CAR-T cells into the same tumor-bearing mice, indicating that c-Jun overexpression enhances preservation of this critical stem-like subset in a cell-intrinsic manner. Although the mechanisms by which c-Jun promotes differentiation of stem/memory-like Tcf1^+^ subsets are unclear, some studies suggest that c-Jun can directly bind to the *Tcf7* locus and disrupt binding of inhibitory AP-1 TFs like IRF4, which promote CD8^+^ T cell exhaustion and impair memory development during chronic infection.^65,106^ Interestingly, endogenous Tpex cells also show strong enrichment for c-Jun DNA binding,^41^ and a recent study demonstrated that neuropilin-1 induced c-Jun down-regulation in endogenous tumor-specific T cells that correlated with poor Tpex self-renewal, terminal differentiation, and resistance to ICI.^107^ These data suggest that c-Jun levels may broadly correlate with sensitivity to ICI among both endogenous and engineered T cells.^65,108–111^

Since the initial study by Lynn et al. showing that c-Jun overexpression renders CAR-T cells “exhaustion resistant” in xenograft models, there has been limited demonstration that c-Jun overexpression is sufficient to enhance CAR-T activity in patients or more clinically relevant mouse models, and the mechanisms limiting c-Jun activity *in vivo* have not been defined.^65,66,112^ Indeed, preliminary data from a phase 1 clinical trial evaluating ROR1 CAR-T cells overexpressing c-Jun indicate that c-Jun overexpression increases the anti-tumor activity and stem-like properties of ROR1 CAR-T cells in patients with ROR1^+^ TNBC and NSCLC but is insufficient to mediate complete responses, mirroring our results in the KP^ROR1^ model.^113^ Our data suggest that PD-1/PD-L1-dependent downregulation of c-Jun is a major factor limiting efficacy of c-Jun.ROR1 CAR-T cells, and this down-regulation may be more pronounced in tumors heavily infiltrated by PD-L1^+^ myeloid cells. Indeed, our data show that, similar to human NSCLC and in contrast to xenograft models, macrophages and neutrophils comprise the dominant immune infiltrate within KP^ROR1^ tumors, highly express PD-L1, and co-localize closely with PD-1^+^ CAR-T cells.^114,115^ Thus, myeloid-derived PD-L1 could contribute to increased PD-1 signaling, c-Jun down-regulation, and ultimate cJun.ROR1 CAR-T dysfunction that is observed in immune-competent models but not xenograft models, which lack PD-L1^+^ myeloid cells.^116^ Although the mechanisms by which the PD-1/PD-L1 axis down-regulates c-Jun in CAR-Ts is unclear, our data indicate that c-Jun down-regulation depends partially on intrinsic PD-1 signaling within CAR-T cells. PD-1 signaling may inhibit c-Jun phosphorylation at sites that prevent binding to E3 ubiquitin ligases, resulting in increased ubiquitin-mediated degradation.^92–97^ In fact, PD-1 signaling has been shown to activate ubiquitin ligases that facilitate degradation of critical components of the TCR and IL-2 signaling pathways, driving T cell dysfunction.^117–119^ Altogether, our data reveal that PD-L1 blockade significantly restores c-Jun levels and cJun.ROR1 CAR-T cell activity *in vivo*, providing a readily translatable strategy to maximize the benefits of c-Jun overexpression in patients.

The KP model is notoriously aggressive, and achieving nearly complete elimination of ROR1^+^ tumor with a single infusion of CAR-T cells and only two weeks of anti-PD-L1 treatment is a major feat. However, our data also indicate that efficacy is likely to be limited by progression of ROR1^null^ tumor. As most targets for CAR-T cells in solid tumors are heterogeneously expressed, escape of antigen^null^ tumor is likely to impair efficacy of even the most potent CAR-T cell products. One limitation of the KP^ROR1^ model is that the ROR1 expressed within tumors is truncated and does not signal. As ROR1 has been shown to play a functional role in promoting tumor proliferation, growth, and metastasis,^19,120–128^ there may be a greater selective pressure against escape of ROR1^null^ tumors in patients that is not appropriately modeled in KP^ROR1^ mice. Nevertheless, strategies to control escape of antigen^null^ tumors will be critical for efficacy of CAR-T therapy in patients with heterogeneous solid tumors. One approach is to engage the endogenous anti-tumor immune response, which may be polyclonal and capable of controlling escape of tumors that lack or downregulate expression of the CAR target. Our studies show that combination of c-Jun CAR-T cells with PD-L1 blockade synergistically boosted numbers of endogenous PD-1^+^ CD8^+^ T cells within KP tumors, though whether these cells contribute to tumor control is not known. As KP tumors lack the high mutational burden of human NSCLC tumors,^129^ additional studies introducing model neoantigens into KP tumors may be needed to determine whether combination of c-Jun CAR-T cells with PD-L1 blockade can synergistically activate endogenous tumor-specific immune responses to control escape of CAR antigen^null^ tumors.

Finally, while c-Jun overexpression significantly enhances CAR-T cell function and response to ICI, it is possible that enhanced CAR-T cell activity may reveal new toxicities *in vivo*. ROR1, like most target antigens for CAR-T cells in solid tumors, shares some expression in normal tissues, including parathyroid, pancreatic islets, and gastric mucosa.^19^ As the CAR used in our study targets human ROR1 in KP tumors and does not cross-react with murine ROR1, we are unable to evaluate the effects of c-Jun CAR-T therapy and PD-L1 blockade on on-target off-tumor toxicity in this model.^21,22^ Although ROR1 CAR-T cells did not induce dose-limiting toxicities in patients with ROR1^+^ CLL, TNBC or NSCLC, 67% of patients experienced reversible grade 3 or 4 pulmonary adverse events, suggesting that lung tissue may be susceptible to toxicity.^23^ Notably, despite the massive accumulation of c-Jun.ROR1 CAR-T cells in the lung upon PD-L1 blockade, we did not observe any toxicity in the KP^ROR1^ model, suggesting that efficacy and safety can be balanced in the absence of CAR target expression on normal tissue. Strategies to enhance CAR-T cell activity, thus, may need to be coupled with logic-gating strategies to restrict CAR-T cell activity to tumor cells to enhance anti-tumor activity without compromising safety. Alternatively, antigens with heterogeneous but tumor-specific expression, like EGFRvIII or the Tn-Muc1 glycoform,^130,131^ may be attractive candidates to target in combination with c-Jun overexpression and/or PD-L1 blockade, as our data show that this treatment strategy significantly reduced tumor burden without toxicity despite ROR1 being expressed in only ∼60% of tumor cells prior to CAR-T infusion.

In summary, our data demonstrate that c-Jun overexpression enables ROR1 CAR-T cells to establish a critical stem-like reservoir *in vivo* that dramatically sensitizes them to enhancement with PD-L1 blockade, providing a strong rationale for combining c-Jun CAR-T cell therapy with ICI to enhance efficacy in patients. Moreover, given that c-Jun levels correlate broadly with sensitivity of both endogenous and engineered T cells to ICI, our work raises the exciting possibility that c-Jun overexpression could be broadly applied to sensitize not only CAR-T, but also TCR-T and tumor-infiltrating lymphocyte (**TIL**) adoptive cell therapies, to enhancement with ICI. Such strategies to sensitize endogenous and engineered T cells to ICI have the potential to extend the transformative benefits of ICI to the majority of cancer patients who remain refractory to ICI.

## MATERIALS AND METHODS

### EXPERIMENTAL MODEL AND SUBJECT DETAILS

#### Animals

C57BL/6 (**B6**) and B6.SJL (**CD45.1**) mice were purchased from Jackson Laboratory. *Kras^LSL-G12D/+^;p53^fl/fl^* (**KP**) mice were generously provided by A. McGarry Houghton (FHCC, Seattle WA). For studies with KP mice, 8-18 week old age-matched and sex-matched mice were used. For all other studies, 8-12 week old age-matched and sex-matched mice were used. Mice of the same sex were randomly assigned to experimental groups or were assigned based on tumor burden such that all experimental groups had similar average tumor volume prior to treatment. All mice were housed and bred at the FHCC (Seattle, WA). All experiments were performed in accordance within the guidelines of the FHCC Institutional Animal Care and Use Committee (PROTO202000031).

#### Cell Lines

The KP1233 (“KP”) tumor and 3TZ “GreenGo” cell lines were generously provided by Tyler Jacks (Massachusetts Institute of Technology). KP-ROR1 tumor cell lines were previously described^21^ and were generated by retroviral transduction of the KP tumor cell line with full-length human ROR1 (UniProt: Q01973) and subsequent FACS sorting of ROR1^+^ cells to >95% purity. All cells were tested bi-monthly for the absence of mycoplasma. KP, KP-ROR1, and 3TZ cell lines were maintained in complete DMEM (DMEM with 10% FBS, 2mM L-glutamine, 100U/ml penicillin/streptomycin, 25mM HEPES). Lenti-X cells for lentiviral packaging were purchased from Clontech. Plat-E cells for retroviral packaging were purchased from Cell Biolabs. All cells were tested bi-monthly for the absence of mycoplasma. KP, KP-ROR1, 3TZ, Plat-E, and LentiX cell lines were maintained in complete DMEM (DMEM with 10% FBS, 2 mM L-glutamine, 100 U/ml penicillin/streptomycin, 25 mM HEPES).

### METHOD DETAILS

#### Cloning of Murine CAR Constructs

The MP71 retroviral vector, which was a gift from W. Uckert (Max Delbruck Center for Molecular Medicine), was modified to encode a human ROR1-specific CAR co-expressed with a truncated murine CD19 transduction marker (**tCD19**). In some experiments, the tCD19 transduction marker was fused to eGFP (tCD19/GFP). The ROR1 CAR possessed a murine CD8α signal peptide (UniProt: P01731, aa1-27), 2A2 scFv^21,132^, human IgG4 short spacer^133^, murine CD28 transmembrane (UniProt: P31041, aa151-177), murine 4-1BB signaling domain (UniProt: P20334, aa211-256) or murine CD28 signaling domain with an LLàGG substitution located at positions 184-185 of the native CD28^134^ (UniProt: P31041, aa178-218), murine CD3ζ (UniProt: P24161, aa52-164), and was linked by a P2A ribosomal skip element to tCD19/GFP. To generate ROR1 CARs co-expressing c-Jun or BATF, the vector was further modified to express murine c-Jun (Uniprot: P05627) or murine BATF (Uniprot: O35284) upstream and linked by a P2A skip element to ROR1 CAR-P2A-tCD19/GFP. To generate the cJun-P2A-Thy1.1 vector, murine c-Jun (Uniprot: P05627) was linked by a P2A skip element to Thy1.1 (Uniprot: P01831). Vectors expressing only the tCD19 or tCD19/GFP marker was used to generate control T cells.

#### Generation of Murine CAR-T Cells

Retrovirus was produced by transient transfection of Plat-E cells with the indicated MP71 vectors. Viral supernatant was harvested 48 hr after transfection, filtered through a 0.45 μm pore filter, and snap-frozen in liquid nitrogen for long-term storage. Cell suspensions were prepared from spleen and peripheral lymph nodes of donor mice by tissue disruption with glass slides and filtered through a 40 μm filter. Murine CD8^+^ and CD4^+^ T cells were enriched from spleens and peripheral lymph nodes of donor mice using untouched negative isolation kits (Stem Cell) and stimulated with anti-CD3/28 Mouse T-Activator Dynabeads (ThermoFisher) at a bead to cell ratio of 1:1 for 24 hr in a 37°C, 5% CO_2_ incubator in complete RPMI (RPM1 1640, 10% heat inactivated FBS, 1 mM sodium pyruvate, 1 mM HEPES, 100 U/ml penicillin/streptomycin, 50 μM β-mercaptoethanol) supplemented with 50 U/ml recombinant human IL-2 (Peprotech). Non-tissue culture plates were coated with 12.5 μg/ml RetroNectin (TaKaRa) according to the manufacturer’s protocol, and plates were loaded with pre-titered virus per well and centrifuged for 2 hr at 3000xg at 32°C. For co-transduction experiments, pre-titered virus was mixed 1:1 and added to pre-coated RetroNectin wells and centrifuged for 2 hr at 3000xg. Murine T cells were harvested and resuspended at 1×10^6^ cells/ml in fresh complete RPMI supplemented with 50 U/ml IL-2. Viral supernatant was aspirated from RetroNectin-coated plates, plates were rinsed with PBS, and T cells were added to each virus-coated well. Plates were then centrifuged at 800g for 30 min at 32°C and returned to 37°C, 5% CO_2_ incubators. After 24 hours, T cells were harvested and resuspended in complete RPMI with 50 U/ml IL-2. T cells were subsequently harvested, counted, and maintained at 1×10^6^ cells/ml in complete RPMI with 50 ng/ml human IL-15 every 1-2 days after. Four days after transduction, magnetic beads were removed and T cell transduction was measured by flow cytometry staining for tCD19, GFP, Thy1.1, ROR1 CAR, and/or c-Jun. Transduced cells were enriched using the EasySep CD19 Positive Selection Kit II (Stem Cell) and either used immediately for *in vitro* or *in vivo* experiments or cryopreserved in complete RPMI supplemented with 10% FBS and 10% DMSO. Cryopreserved cells were thawed in a 37°C water bath, washed, counted, re-confirmed for viability, tCD19, GFP, Thy1.1, ROR1 CAR, and/or c-Jun expression by flow cytometry, and used immediately for experiments.

#### Generation of PD-1^KO^ CAR-T cells

Ribonucleoprotein (**RNP**) complex was prepared by combing PD1-sgRNA (5′-ACAGCCCAAGUGAAUGACCA) and Cas9 protein (Horizon Cat# CAS12206) at a 3:1 molar ratio. Lonza P3 buffer (Catalog #: V4XP-3024) was prepared according to package insert. Murine CD8^+^ T cells were isolated from spleens and peripheral lymph nodes of donor mice and stimulated with anti-CD3/28 Mouse T-Activator Dynabeads (ThermoFisher) at a bead to cell ratio of 1:1 for 24 hr in a 37°C, 5% CO_2_ incubator in complete RPMI supplemented with 50 U/ml recombinant human IL-2, as described above. After 24 hr of stimulation, CD8^+^ T cells were harvested, de-beaded, and resuspended in Lonza P3 buffer and RNP (4×10^6^ cells per 80 ul). The cell/RNP mixture was transferred to a Lonza 4D-Nucleofector® X Kit L cuvette (Catalog #: V4XP-3024) and electroporated in a Lonza 4D Nucleofector Core Unit using the “CA137” program. Following electroporation, cells were rested at 37°C for 15 mins. Mouse T-Activator anti-CD3/28 beads were added back to the cell suspension 1:1 with T-cells. T cells were then added to retrovirus-coated plates and cultured as described above. Five days post-electroporation, genomic DNA (**gDNA**) was extracted from a subset of cells using the QuickExtract™ DNA Extraction Solution (BioScience Technologies, QE0905T). The 900 bp region surrounding the sgRNA was amplified via PCR with forward (5’-CACCTCTAGTTGCCTGTTCT) and reverse (5’-CACCTGTAAAACCCACCTAA) primers. gDNA was sequenced by ACGT DNA Sequencing Services, using the primer sequence CACCTCTAGTTGCCTGTTCT. Indel frequencies were determined using the ICE analysis tool provided by Synthego Corporation. To further validate efficient knockdown of PD-1 expression, T cells were stimulated with 50ng/ml PMA and 1 μg/ml ionomycin in 0.2 ml complete RPMI in 96-well U-bottomed plates (Costar) at 37°C, 5% CO_2_ for 24 hr and analyzed by flow cytometry for expression of PD-1 by flow cytometry.

#### Generation, Titration, and Intratracheal Administration of Cre Lentivirus

We previously described modification of the HIV7 lentiviral vector^135^ to encode Cre recombinase, firefly luciferase (**ffluc**), and truncated human ROR1 (**hROR1t**) (UniProt: Q01973, aa 1-462), with each transgene linked by P2A and T2A ribosomal skip elements (Cre-P2A-ffluc-T2A-hROR1t).^21^ We further modified this vector to reposition hROR1t in the 5’ position, by expressing hROR1t (UniProt: Q01973, aa 1-462) linked by a P2A ribosomal skip element to Cre recombinase (**hROR1-P2A-Cre**). As a control, the HIV7 vector was modified to express Cre only.

Lentivirus was produced by transient calcium phosphate transfection of the packaging cell line LentiX with the indicated HIV7 lentiviral vectors, psPAX2 (Addgene #12260), and VSVg envelope. Viral supernatant was harvested 24, 48 and 72 hr after transfection, filtered through a 0.45-μm pore filter, and stored at 4°C for up to 1 week until ready for ultracentrifugation. Lentivirus was concentrated by mixing filtered lentivirus with 40% polyethylene glycol (PEG, Sigma) at a PEG to virus ratio of 1:3 for 12-24 hr at 4°C. The virus/PEG mixture was then centrifuged at 1500g for 45 min at 4°C, supernatant was aspirated, and the virus pellet was resuspended in 30 ml serum-free DMEM. Lentivirus was further concentrated in an Optima L-90K ultracentrifuge (Beckman Coulter) at 24,500 rpm for 90 min at 4°C. The final virus pellet was resuspended in 0.5 – 1 ml serum-free DMEM by vortexing for 1-3 hr at 4°C, aliquoted, and frozen at –80°C for long-term storage.

Cre lentivirus was titered using 3TZ “Green Go” cells. Briefly, 5×10^4^ 3TZ cells were plated in 1 ml of complete DMEM in 12-well plates. 5-6 hours later when cells were adherent, supernatant was aspirated and media was replaced with 0.5 ml complete DMEM containing serial dilutions of thawed Cre lentivirus (e.g. 10-fold serial dilutions from 1:10 to 1:10,000) and 8 μg/ml polybrene. After 24 hr, media was replaced with 1 ml complete DMEM and cells were passaged for 3-4 days before analysis by flow cytometry for GFP and/or hROR1 expression. Virus titer was calculated according to the following formula: Titer (pfu/μl) = [(5×10^4^) * (%GFP^+^ cells) / 100] / [Volume of virus added (μl)]. Lung tumors were induced in KP mice by intratracheal intubation and inhalation of 2×10^4^ pfu Cre-expressing lentivirus as reported.^62^

#### CAR-T Cell Treatment of KP^ROR1^ mice

KP mice were scanned by micro-computed tomography (micro-CT) ∼10-11 weeks post-infection and one week before T cell infusion, and tumor burden was calculated for each mouse using 3D Slicer image computing platform. Mice were distributed into treatment groups such that the average tumor burden was as similar as possible prior to treatment to control for variability in induction of tumors. For all T cell infusion experiments, KP mice were injected intraperitoneally with 150 mg/kg cyclophosphamide and 6-24 hr later were injected intravenously by retro-orbital injection with 6×10^6^ live tCD19^+^ control or CAR-T cells (1:1 ratio of CD8:CD4 T cells). For co-transfer experiments, ROR1 CAR-T and cJun.ROR1 CAR-T cells were mixed 1:1 prior to infusion. Mice were monitored for weight, body condition score, tumor burden, and survival where indicated. All mice in each experiment were sacrificed when any individual mice showed clinical signs of severe disease or 20 percent weight loss.

#### Lung Metastatic Transplantable KP Tumor Model

5×10^4^ KP-ROR1 tumor cell lines were injected intravenously via tail vein into 8–12 week old B6 mice to induce development of metastatic lung tumors. After 21 days, tumor-bearing mice were injected intraperitoneally with 150 mg/kg cyclophosphamide and 6-24 hr later were injected with 6×10^6^ live tCD19^+^ control or CAR-T cells (1:1 ratio of CD8:CD4 T cells). Mice were monitored for weight, body condition score, tumor burden, and survival where indicated. All mice in each experiment were sacrificed when any individual mice showed clinical signs of severe disease or 20 percent weight loss.

#### Anti-PD-L1 Treatment

In some experiments, mice were treated with 200 ug anti-PD-L1 antibody (BioXcell, clone: 10F.9G2) intraperitoneally every 3-4 days beginning 1 day post T cell transfer and continuing for 14 days (5 total doses).

#### Preparation of Tissues for Flow Cytometry and RNAseq

Cell suspensions were prepared from murine spleen and peripheral lymph nodes by tissue disruption with glass slides and filtering through a 40 μm filter, and lysing with ACK lysis buffer (Gibco). For analysis of lung tumors, mice were injected with 400 ng PECy7-conjugated anti-CD45 antibody (clone 30-F11, BioLegend) intravenously via retro-orbital injection 5 min prior to euthanasia to distinguish vascular and non-vascular cells in the lung. Whole lungs were placed in GentleMACS C Tubes with 6 ml of complete RPMI supplemented with 2 mg/ml collagenase IV (Worthington) and 80 U/ml DNAse I (Worthington). Lungs were minced using the “m_impTumor_01_01” program on a GentleMACS Octo Dissociator and incubated in a 37°C shaker at 120 rpm for 30 min. Cells were filtered through a 100 μm filter, lysed with ACK lysing buffer (Gibco), and resuspended as single cell suspensions for downstream analysis. All flow cytometry analyses of tumor-bearing lungs were performed on live CD45-PECy7^−^ cells. For bulk RNAseq and single cell RNAseq experiments, lung single cell suspensions were cryopreserved in complete RPMI supplemented with 10% FBS and 10% DMSO.

#### Flow Cytometry

Cells were stained using the Live/Dead Fixable Aqua Dead Cell stain kit (Invitrogen) according to the manufacturer’s protocol. For surface staining, cells were incubated at 4°C for 30 min in staining buffer (PBS, 2% FBS) with the following directly conjugated antibodies for murine proteins (from BioLegend unless otherwise specified): anti-CD4 (RM4-5), –CD8 (53-6.7), –CD45.1 (A20), – CD45.2 (104), –CD45 (30-F11), –CD3 (145-2C11), –CD19 (eBio1D3, Thermo Fisher), –PD-1 (29F.1A12), –LAG-3 (C9B7W), –TIM3 (RMT3-23), –Tox (REA473, Miltenyi), –Thy1.1 (OX-7), Thy1.2 (53-2.1), –B220 (RA3-6B2), NKp46 (29A1.4), NK1.1 (PK136), –CD11b (M1/70), –CD11c (N418), –Ly6C (HK1.4, Thermo Fisher), –Ly6G (1A8), –F4/80 (BM8), I-A/I-E (M5/114.15.2, Thermo Fisher), –CD24 (M1/69), –CD64 (X54-5/7.1), –SiglecF (E50-2440, BD Biosciences), –PD-L1 (10F.9G2), –EpCAM (G8.8, Thermo Fisher); or with the following directly conjugated antibodies for human proteins: anti-ROR1 (2A2, Miltenyi Biotec). Recombinant Fc-hROR1 (Fred Hutchinson Cancer Center Protein Core) and anti-human IgG secondary antibody (HP6017, BioLegend) was used to measure ROR1 CAR expression. For intracellular staining, cells were surface stained as described, washed and permeabilized for 20 min with eBioscience Fix/Perm kit at 4°C. Cells were stained for 30 min at 4°C in 1X Perm/Wash buffer (eBioscience) with anti-mouse IFN-γ (XMG1; Thermo Fisher), –TNFα (MP6XT22, Thermo Fisher), –IL-2 (JES6-5H4), –Foxp3 (FJK-16S), –Ki67 (B56, BD Biosciences), –Tcf1 (C63D9, Cell Signaling), and/or –cJun (60A8, Cell Signaling). For intracellular cytokine staining following restimulation, cells were stimulated with 50ng/ml PMA and 1 μg/ml ionomycin in 0.2 ml complete RPMI in 96-well U-bottomed plates (Costar) at 37°C, 5% CO_2_ for 6 hr. GolgiPlug (BD Biosciences) was added to all wells according to the manufacturer’s protocol at the beginning of co-culture. All data were acquired on FACS Symphony or Canto II flow cytometers (BD Biosciences) and analyzed using FlowJo software (Treestar).

#### In Vitro Proliferation Assay

For analysis of CAR-T cell proliferation *in vitro*, cells were incubated with 10 uM Cell Trace Blue (Thermo Fisher) in PBS for 9 mins at 37°C and washed with 100% FBS. Cell Trace Blue-labeled T cells were stimulated with 50ng/ml PMA and 1 μg/ml ionomycin in 0.2 ml complete RPMI in 96-well U-bottomed plates (Costar) at 37°C, 5% CO_2_ for 72 hr and analyzed by flow cytometry. Proliferation index was calculated from Cell Trace Blue dye dilution profiles using FlowJo software (Tree Star).

#### Immunohistochemistry

Lungs were fixed in 10% neutral-buffered formalin. Formalin-fixed paraffin-embedded tissues were sectioned at 4 microns onto positively-charged slides and baked for 1 hr at 60°C. The slides were then dewaxed and stained on a Leica BOND Rx autostainer (Leica, Buffalo Grove, IL) using Leica Bond reagents for dewaxing (Dewax Solution). Antigen retrieval was performed at 100°C for 20 min using Leica Epitope Retrieval Solution 1 (Leica AR9961). Endogenous peroxidase was blocked with H2O2 (in the Leica kit D59800) for 5 min followed by protein blocking with TCT buffer (0.05 M Tris, 0.15 M NaCl, 0.25% Casein, 0.1% Tween 20, pH 7.6 +/− 0.1) for 10 min. Primary rabbit anti-ROR1 antibody at 1:500 (Cell Signaling #16540) or rabbit anti-CD8 antibody at 1:200 (Cell Signaling #98941) was applied for 60 minutes followed by the Refine reagent Rbt-HRP polymer (in the Leica kit D59800) for 12 minutes. Staining was visualized using Mixed Refine DAB (in the Leica kit D59800) and then counter-stained with Hematoxylin (in the Leica kit D59800) for 4 minutes, rinsed and dehydrated and then cover-slipped. Percent ROR1^+^ cells and CD8^+^ localization within tumors was quantified using HALO software (Indica Labs).

#### Single-Cell RNA Sequencing

Cryopreserved lung tumor cell suspensions and CAR-T infusion products were thawed, washed, and labeled with TotalSeq-C anti-mouse Hashtags (Biolegend) according to the manufacturer’s protocol. Following hashing, equal numbers of live cells were pooled from 3 mice in each treatment group to form 4 pools: D9 ROR1 CAR, D9 cJun.ROR1 CAR, D30 ROR1 CAR, D30 cJun.ROR1 CAR. For analysis of T cells, CD8^+^ T cells were isolated from each pool using the Mouse CD8a Positive Selection Kit II (Stem Cell). For analysis of the TME, each pool was stained Live/Dead Fixable Aqua Dead Cell stain kit (Invitrogen) and live CD45-PECy7^−^ cells were sorted. For analysis of CAR-T infusion products, CAR-T cells were used directly after hashing. Libraries from CAR-T infusion product, T cell, and TME samples were prepared using the 10X Genomics Chromium GEM-X Single Cell 5’ Kit as per the manufacturer’s instructions and were sequenced on an Illumina NextSeq 2000 according to the standard 10X Genomics protocol.

Resulting reads were aligned to modified reference genome which includes sequences for the ROR1 scFv, tCd19-GFP, and cJun transgenes, using CellRanger (v.8.0.0) software from 10X Genomics. HDF5 matrix files were imported into R using the Seurat v5 package. Cells whose UMI counts for mitochondrial genes greater than 10% were excluded from downstream analysis. Cells with between 200 and 7000 genes expressed per cell were included in the analysis. Hashtag oligos (HTO) assay was demultiplexed using Seurat function, HTODemux, to assign hashes and identify cells with more than one hash to be excluded. Scrublet was used to identify potential multiplets which were excluded from downstream analysis. We obtained the following total numbers of cells for each sample after filtering cells for quality: 25458 cells (T cells) and 39639 cells (TME).

#### Clustering, Differential Expression, Geneset Scores, and Differential Abundance Analysis

Clustering was performed using Seurat on CAR+ cell clusters in the T cells population, numbering 8766 cells. For unsupervised clustering of cell expression profiles, expression matrices were first normalized using SCTransform, while regressing out mitochondrial percentage. We then performed PCA dimension reduction, followed by FindNeighbors, FindClusters with resolution of 0.8, and UMAP reduction on the first 10 dimensions. Cell annotation was done using cluster markers from FindAllMarkers on upregulated genes using the following settings: log fold change threshold of 0.25, p-adjusted threshold of 0.05, and genes expressed in at least 10% of the cluster (mitochondrial and ribosomal genes were not included in input features). Relative expression of specific gene sets across cells were calculated using AddModuleScore function.

Differential abundance analysis was performed using miloR package (v 2.0.0) to identify enrichment of either treatment group (ROR1 CAR-T versus cJun.ROR1 CAR-T). K-nearest neighbor (k-NN) graph was constructed using miloR standard workflow, using k = 30 and prop = 0.5 to define neighborhoods. We included “sample_hash” variable as covariate to account for variation across different mouse IDs and time points. Enrichment was assessed using Spatial false discovery rate (SpatialFDR) threshold of < 0.05. Each neighborhood was annotated based on the predominant cell type, with those containing less than 70% of a single cell type labeled as “Mixed” neighborhoods. We used a set of 1000 highly variable genes, identified with modelGeneVar() and getTopHVGs() from scran R package (v.1.32.0), as input features for differential gene expression analysis between all neighborhoods enriched in ROR1 CAR-T versus cJun.ROR1 CAR-T. Neighborhood expression profiles were aggregated by treatment group, and significant markers were identified using adjusted p-value < 0.05.

#### Xenium Custom Probe Panel Design and In Situ Gene Expression

A 100 gene custom add on *in situ* gene expression panel was generated with the Xenium Panel Designer Tool by 10x Genomics (online tool). The custom design was informed by annotated 5’ single-cell RNA sequencing data described in this study (Fig.3C) and included custom probes for the ROR1-Cre (*SS109-CRtumor*) and ROR1 CAR (*ROR1_CAR-*T) transgenes, to enable detection of ROR1^+^ tumor and CAR^+^ T cells. Probes for 99 genes identified as differentially expressed and biologically informative from the combined dataset were developed to augment the Xenium Mouse Tissue Atlasing Panel, which includes 379 genes for identifying major mouse cell types. Throughout the panel design process, we prioritized probe coverage for key cell types of interest, ensuring robust representation for populations such as macrophages,

B cells, dendritic cells, fibroblasts, neutrophils, NK cells, various T cell subsets (including Th17 and Treg), and multiple tumor/stroma subpopulations. Probe selection was optimized to maximize detection sensitivity across these cell types while carefully managing allocation of the detection budget to minimize optical crowding. Each gene’s probe sets were chosen to balance coverage and specificity, accommodating the biological heterogeneity of the sample.

Formalin-fixed paraffin embedded (FFPE) tissue blocks were sectioned and placed on the Xenium slides as outlined in the 10X Genomics Xenium In Situ Protocols FFPE Tissue Preparation Guide CG000578 Rev. C. Slides were incubated in a covered container in the Eppendorf Innova at 42°C for 3 hr to further adhere the tissue sections to the slide. Slides were then stored in a desiccator at room temperature in the dark until ready to proceed. Deparaffinization and decrosslinking were performed per the 10X Genomics CG000580-Rev C protocol. Briefly, slides were incubated in a covered container in the Eppendorf Innova at 60°C for 2 hr. After cooling to room temperature, the tissue sections underwent a series of submersions in xylene, ethanol, and nuclease free water to complete the deparaffinization process. The slides were loaded into the provided Xenium cassettes, a decrosslinking buffer added, and the slides incubated on the thermocycler adapter to decrosslink and permeabilize the sections. Xenium In Situ Gene Expression User Guide (CG000584 Rev. F) was followed for subsequent steps as follows. Probe preparation was performed by heating the Mouse Multi-Tissue panel + custom add on (described above) to 95°C for 2 min before snap cooling on ice for 1 min. The mix of probes and buffer was added to the slides and incubated overnight for a minimum of 16 hr at 50°C. Slides were washed with PBS-T and incubated with a post hybridization wash buffer for 30 min at 37°C. The padlock probes were then ligated at the junction by incubating with a ligase enzyme for 2 hr at 37°C. Following ligation and washing with PBS-T, rolling circle amplification was performed for 2 hr at 30°C\ to amplify the signal from each probe. Slides were prepared for imaging by applying an autofluorescence quenching solution followed by nuclei staining before loading on to the Xenium instrument as instructed by Xenium Analyzer User Guide (CG000584 Rev. E). Upon completion of the run, slides underwent H&E staining following quencher removal in a 10 mM sodium hydrosulfite solution for 10 min. Staining was performed according to Xenium In Situ Gene Expression-Post-Xenium Analyzer H&E staining demonstrated protocol (CG000613 Rev. B).

#### Xenium Analysis

Raw Xenium spatial transcriptomic data was segmented using the Proseg algorithm.^136^ All subsequent data processing and analysis was performed in Python using the scanpy v1.9.8^137^ and squidpy v1.5.0^138^ packages. For each sample, low quality cells with fewer than 10 total transcripts were removed, and genes detected in fewer than 5 cells were removed. Transcriptional data was then log-normalized and PCA-transformed. A nearest-neighbor graph was constructed and used to generate UMAP representation and Leiden clusters. CAR-T cell clusters were identified based on expression of the *ROR1_CAR-T* marker. Those CAR-T cells were then separately subclustered: a nearest-neighbors graph was constructed based on expression of only T cell marker genes (*Tcf7, Mki67, Pdcd1, Havcr2, Ifng, Gzmb, Prdm1, Tox, Foxp3, Entpd1*) for identification of relevant T cell subsets, which was then used for generation of a UMAP and Leiden subclusters.

To identify spatial regions (tumor, airway/blood vessel, and normal lung parenchyma), an additional nearest-neighbor graph was constructed based on spatial coordinates of cells. A joint graph was constructed from the two nearest-neighbor graphs (the one constructed from spatial coordinates as well as the graph constructed from gene expression), with the spatial coordinate graph weighted at between 0.5 and 0.6, depending on the sample. Leiden clustering was then performed using this joint graph, with resolution 0.7. Resulting spatial clusters were combined in some cases, using the H&E image as a reference. To identify individual tumors, Leiden clustering based on only the spatial coordinate nearest-neighbor graph was performed on the cells identified as belonging to tumor regions. Clusters were again combined when necessary, according to the H&E image. Abundance analysis was performed by calculating the relative proportion of each cluster of interest out of the total number of cells in the region of interest (individual tumors or entire non-tumor region). Proximity analysis was performed using the co-occurrence and neighborhood enrichment functions in squidpy.

## QUANTIFICATION AND STATISTICAL ANALYSIS

All data are presented as the mean values ± SEM. Statistical significance was determined by one-way ANOVA with Tukey’s post-test, two-way ANOVA with Tukey’s post-test, log-rank Mantel-Cox test, unpaired Student’s two-way t-test, or paired Student’s two-way t-test as indicated in figure legends using Prism software (Graphpad). Statistical significance was established at the levels of *, p<0.05; **, p<0.005; ***, p<0.0005; ****, p<0.0001.

### List of Supplementary Materials

Fig. S1 to S6.

## Acknowledgements

We wish to thank Ekram Gad, Elena Carlson, and Tracy Goodpaster in Shared Resources and David Sowerby in the Fred Hutch Innovation Lab for technical services, as well as Joshua Veatch, Mark Headley, and Stan Riddell for helpful discussions and reagents. Treatment schematic figures were created with BioRender. This research was also supported by the Preclinical Imaging, Comparative Medicine, and Experimental Histopathology Shared Resources of the Fred Hutch/University of Washington Cancer Consortium.

## Funding

National Cancer Institute, R37CA276285, (S.S.)

American Lung Association, Lung Cancer Discovery Award (S.S.)

V Foundation, Bob Bast Translational Research Grant (S.S.)

Cancer Center Support Grant, P30 CA015704 (S.S.)

Lyell Immunopharma, Sponsored Research Agreement (S.S.)

National Institutes of Health, T32 5T32GM007266-46 (A.J.S.).

## Author Contributions

A.J.S and S.S. conceived the study, designed experiments, and wrote the manuscript. A.J.S. performed experiments, analyzed the data, and interpreted results. C.S., A.R.W., A.L., A.E., and E.N. generated and performed analysis of spatial transcriptomic data. T.H., S.B., E.F., X.W., S.P, and S.F. performed analysis of single cell RNA-seq data. M.G.K, S.O., W.S.N., S.G., E.B., V.Z., and M.S. performed experiments and analyzed data.

## Data and Materials Availability

Xenium spatial transcriptomic data are deposited in Zenodo and can be accessed under DOI: 10.5281/zenodo.14775896 (or at zenodo.org/records/14775896). Single cell RNAseq (scRNAseq) data are deposited in the Gene Expression Omnibus under the accession number GSE288687. Further information and requests for resources and reagents should be directed to and will be fulfilled by the Corresponding Author, Shivani Srivastava.

## Supplemental Information

**Supplementary Fig. 1. Modification of the lentiviral hROR1-Cre vector improves co-expression of hROR1 and Cre recombinase.** A) Schematic of lentiviral constructs used to induce tumors in KP mice. ffluc = firefly luciferase. hROR1t = truncated human ROR1. Cre = Cre recombinase. B) Flow cytometry plots showing expression of hROR1 in GFP^+^ 3TZ^LSL-GFP^ “GreenGo” Cre-reporter cells infected with the indicated lentiviruses. C) Representative IHC staining (left) and summary (right) of ROR1 expression on lung tumors in KP mice infected with Cre or hROR1-Cre lentivirus. Data are representative of 3-5 independent experiments.

**Supplementary Fig. 2. Overexpression of AP-1 family TFs does not substantially alter ROR1 CAR-T cell function prior to infusion**. A) Flow cytometry plots showing ROR1 CAR, c-Jun, and BATF expression in CD8^+^ T cells untransduced or transduced with ROR1 CAR, cJun.ROR1 CAR, or BATF.ROR1 CAR vectors prior to infusion. B) Flow cytometry plot showing Tcf1 expression in CAR-T cell infusion products relative to ROR1 CAR-T cells within KP^ROR1^ tumors 8 days post-infusion (grey). C) Proliferation of CAR-T cells 72 hours post-stimulation with PMA and ionomycin *in vitro*. N=3 technical replicates. Two-way ANOVA with Tukey’s post-test. D) Representative plot showing dilution of Cell Trace Blue dye upon stimulation with PMA and ionomycin in CD8^+^GFP^+^ CAR-T cell infusion product. E) Intracellular cytokine staining analysis of CD8^+^GFP^+^ CAR-T cell infusion products upon stimulation with PMA and ionomycin *in vitro*. One-way ANOVA with Tukey’s post-test. Data are representative of 3 independent experiments.

**Supplementary Fig. 3. Sub-clustering of CAR-T cells from single cell RNA-seq data.** A) Unsupervised clustering (left) and *Cd3e* expression (right) of CAR-T infusion product and CD8^+^ cells enriched from KP^ROR1^ tumors 9 and 30 days post-CAR-T infusion. B) Expression of *Rag1* among clusters in (A). C) Coloring of clusters in (A) by sample hash. N = 3 mice per group. D) Sub-clustering of cells expressing *Cd3e* and excluding *Rag1* (left) and expression of *Cd3e* (middle) and *CAR* transgene (right) among T cell subclusters. E) Sub-clustering of *CAR^+^* clusters (left) and expression of *CAR* among CAR-T subclusters (right). F, G) Enrichment of various gene sets for memory/stem-like T cells (F) or terminally exhausted T cells (G) among CAR-T subclusters.

**Supplementary Fig. 4. c-Jun overexpression improves activity of both ROR1.BBζ and ROR1.28ζ CAR-T cells, but tumors ultimately progress.** A) Frequency of GFP^+^ CAR-T cells of CD8^+^ T cells in lungs 8 days post-infusion into B6 mice bearing transplanted KP-ROR1 lung tumors. N=3-4 mice per group. One-way ANOVA with Tukey’s post-test. B, C) Inhibitory receptor expression (B) and cytokine production upon *ex vivo* restimulation with PMA and ionomycin (C) in CD8^+^GFP^+^ CAR-T cells in lungs 8 days post-infusion into B6 mice bearing transplanted KP^ROR1^ lung tumors. N=3-4 mice per group. One-way ANOVA with Tukey’s post-test. D) Survival of B6 mice bearing transplanted KP^ROR1^ tumors and treated with the indicated T cell products. N=4 mice per group. Log-rank Mantel-Cox test. Data are representative of 2 independent experiments.

**Supplementary Fig. 5. Annotation of TME single cell RNAseq clusters.** A) Unsupervised clustering of live cells sorted from KP^ROR1^ tumors of mice left untreated, or 9 or 30 days post-infusion with ROR1 CAR-T or cJun.ROR1 CAR-T cells. B, C) Clusters in (A) colored by time point (B) or treatment group (C). D) Dot plot showing expression of various lineage-defining genes across clusters in (A). E,F) UMAP (E) and violin plot (F) showing expression of *Cd274* (encodes PD-L1) across clusters in (A). G) Expression of PD-L1 on CD45^−^EpCAM^+^ROR1^+^ KP tumor cells, CD11c^+^F4/80^+^ macrophages, and CD11b^+^Ly6G^+^ neutrophils within KP^ROR1^ lung tumors left untreated (black) or 9 days post-infusion of ROR1 CAR-Ts (pink) or cJun.ROR1 CAR-Ts (teal). N=3-5 mice per group. One-way ANOVA with Tukey’s post-test.

**Supplementary Fig. 6. Annotation of 10X Xenium clusters.** Lungs from KP^ROR1^ mice were harvested 7 days post-treatment with ROR1 CAR-Ts + vehicle, ROR1 CAR-Ts + anti-PD-L1, cJun.ROR1 CAR-Ts + vehicle, or cJun.ROR1 CAR-Ts + anti-PD-L1 and analyzed by 10X Xenium. Dot plot summarizes expression of various genes defining each cluster.

