## Supplemental Figures S1-S6 for "Modulating AP-1 enables CAR-T cells to establish an intratumoral PD-1^+^Tcf1^+^ stem-like reservoir and overcomes resistance to PD-1 axis blockade"

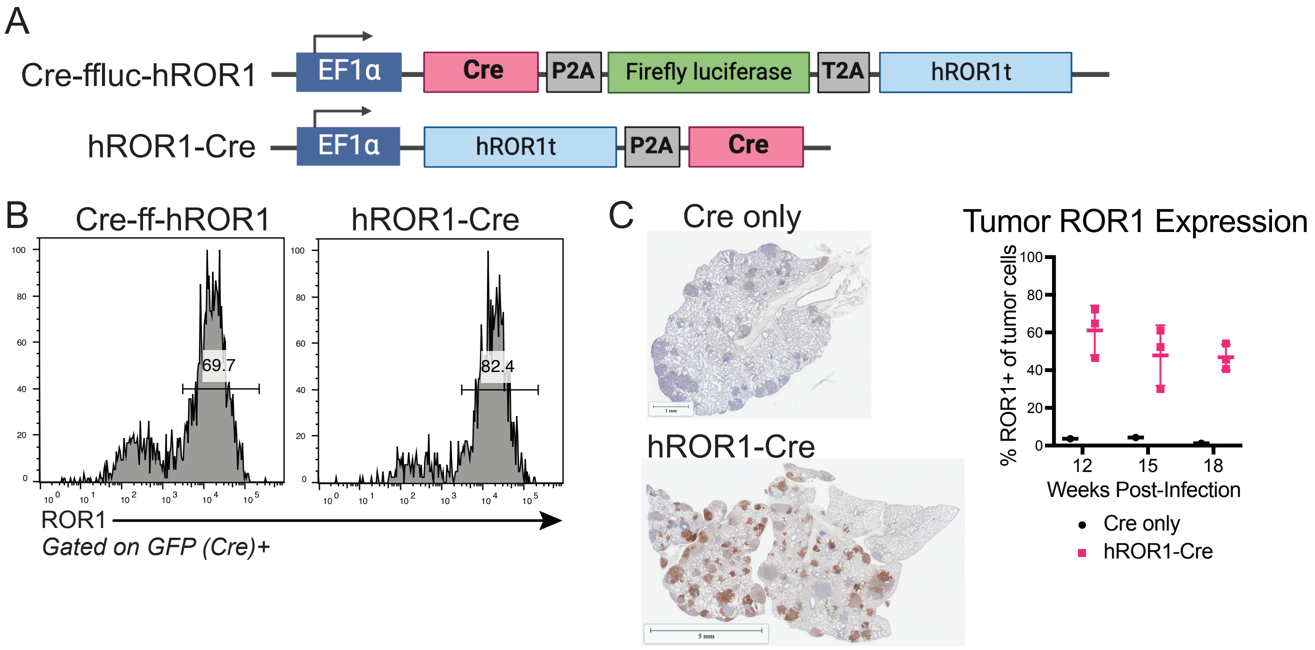


**Figure S1. Modification of the lentiviral hROR1-Cre vector improves co-expression of hROR1 and Cre recombinase.** A) Schematic of lentiviral constructs used to induce tumors in KP mice. ffluc = firefly luciferase. hROR1t = truncated human ROR1. Cre = Cre recombinase. B) Flow cytometry plots showing expression of hROR1 in GFP+ 3TZLSL-GFP "GreenGo" Cre-reporter cells infected with the indicated lentiviruses. C) Representative IHC staining (left) and summary (right) of ROR1 expression on lung tumors in KP mice infected with Cre or hROR1-Cre lentivirus. Data are representative of 3-5 independent experiments.

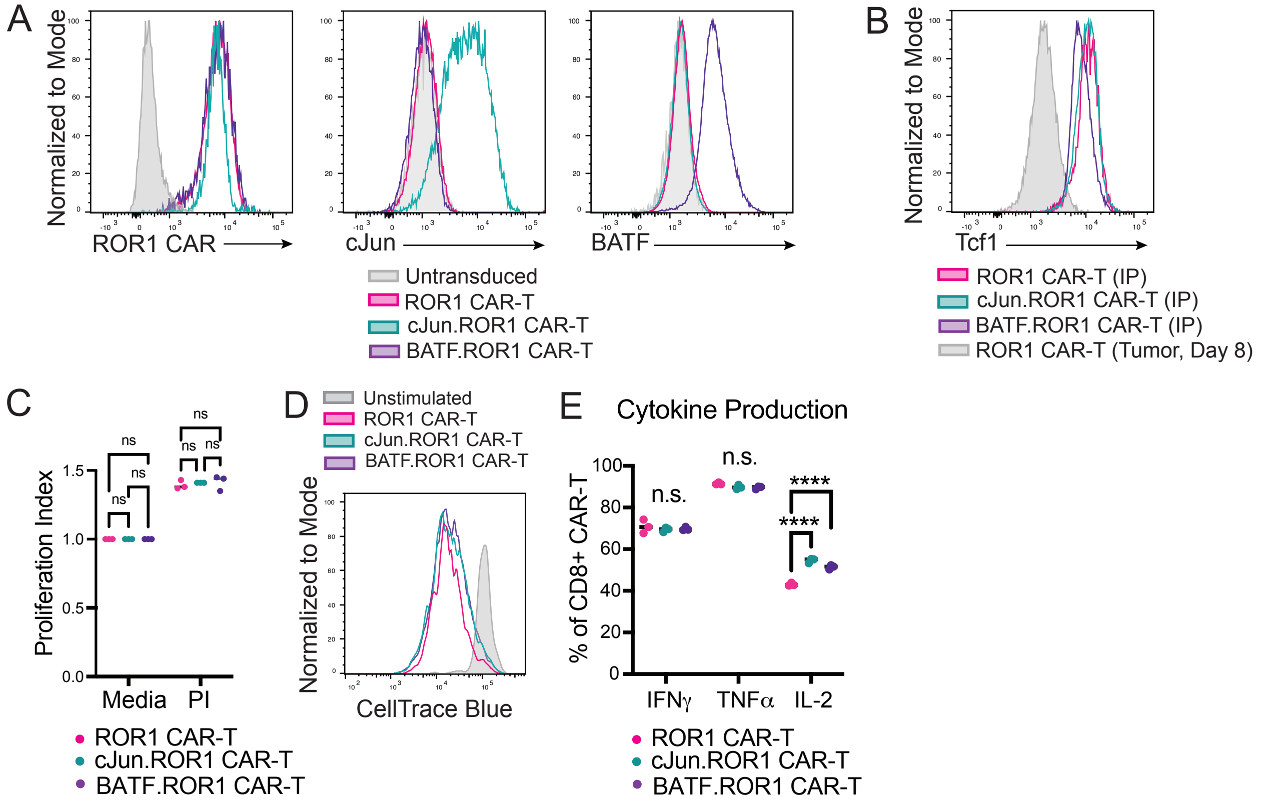


**Figure S2. Ooverexpression of AP-1 family TFs does not substantially alter ROR1 CAR-T cell function prior to infusion.** A) Flow cytometry plots showing ROR1 CAR, c-Jun, and BATF expression in CD8+ T cells untransduced or transduced with ROR1 CAR, cJun.ROR1 CAR, or BATF.ROR1 CAR vectors prior to infusion. B) Flow cytometry plot showing Tcf1 expression in CAR-T cell infusion products relative to ROR1 CAR-T cells within KP-ROR1 tumors 8 days post-infusion (grey). C) Proliferation of CAR-T cells 72 hours post-stimulation with PMA and ionomycin in vitro. N=3 technical replicates. Two-way ANOVA with Tukey's post-test. D) Representative plot showing dilution of Cell Trace Blue dye upon stimulation with PMA and ionomycin in CD8+GFP+ CAR-T cell infusion product. E) Intracellular cytokine staining analysis of CD8+GFP+ CAR-T cell infusion products upon stimulation with PMA and ionomycin in vitro. One-way ANOVA with Tukey's post-test. Data are representative of 3 independent experiments.

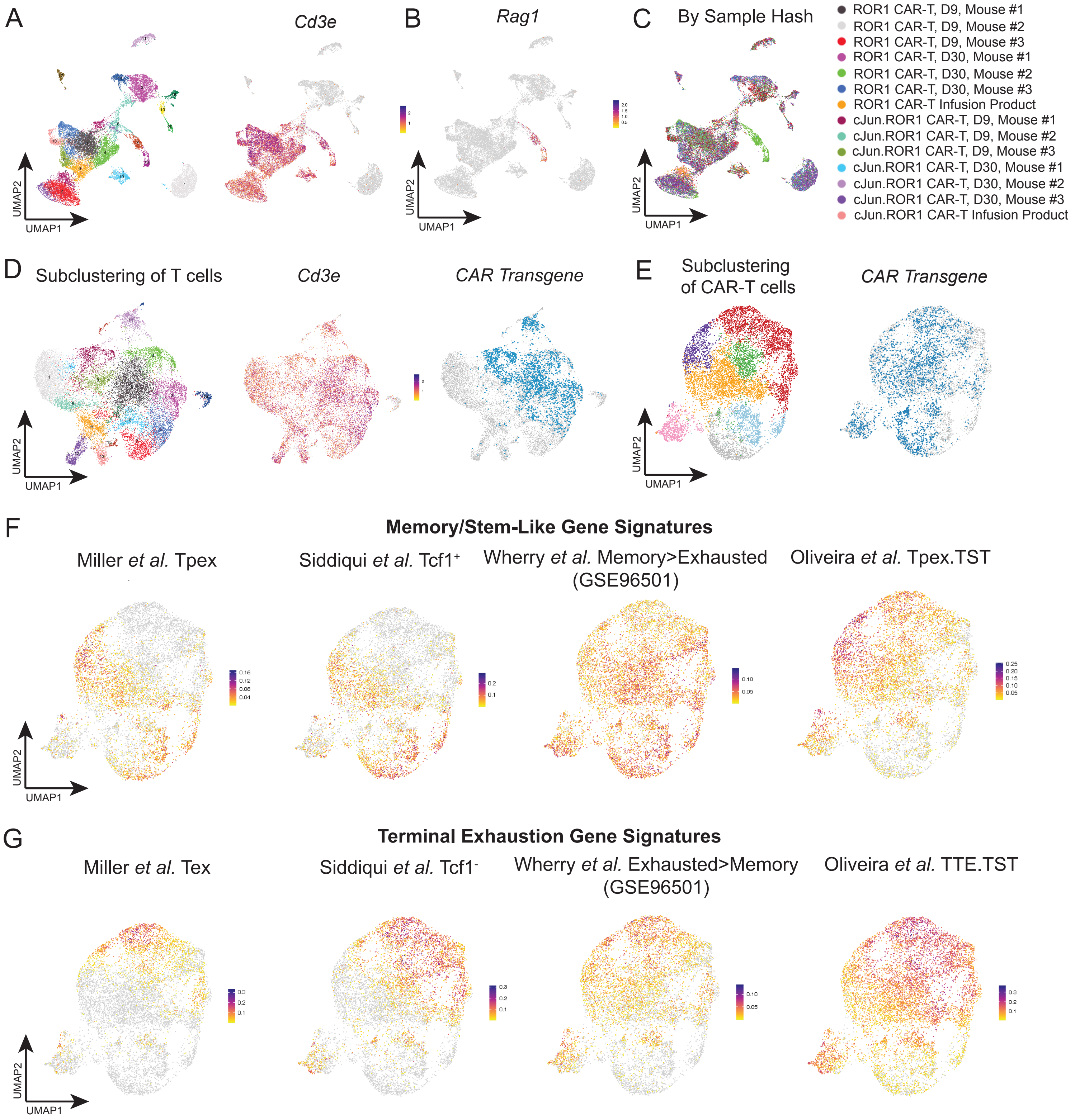


**Figure S3. Sub-clustering of CAR-T cells from single cell RNA-seq data.** A) Unsupervised clustering (left) and *Cd3e* expression (right) of CAR-T infusion product and CD8+ cells enriched from $KP^{ROR1}$ tumors 9 and 30 days post-CAR-T infusion. B) Expression of *Rag1* among clusters in (A). C) Coloring of clusters in (A) by sample hash. N = 3 mice per group. D) Sub-clustering of cells expressing *Cd3e* and excluding *Rag1* (left) and expression of *Cd3e* (middle) and *CAR* transgene (right) among T cell subclusters. E) Sub-clustering of *CAR*+ clusters (left) and expression of CAR among CAR-T subclusters (right). F, G) Enrichment of various gene sets for memory/stem-like T cells (F) or terminally exhausted T cells (G) among CAR-T subclusters.

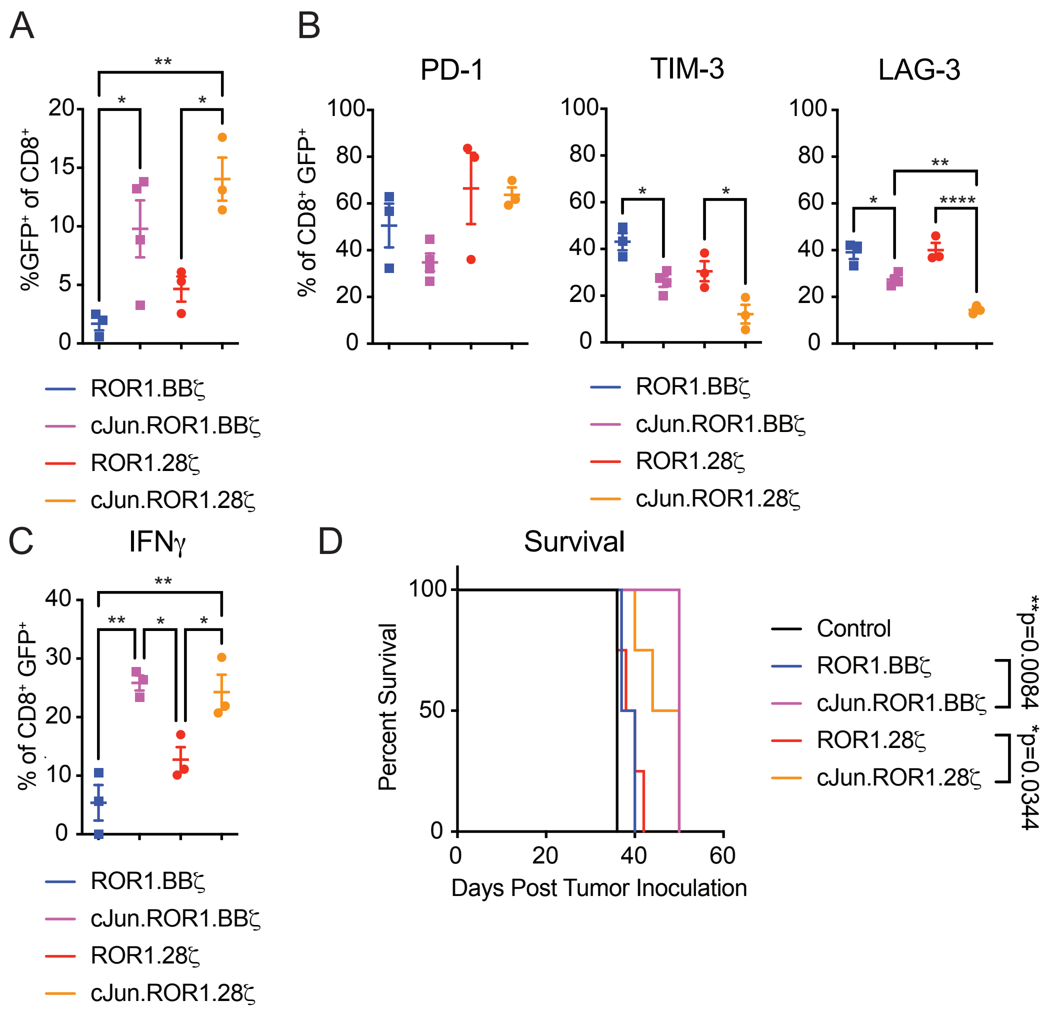


**Figure S4. c-Jun overexpression improves activity of both ROR1.BBζ and ROR1.28ζ CAR-T cells, but tumors ultimately progress.** A) Frequency of GFP+ CAR-T cells of CD8+ T cells in lungs 8 days post-infusion into B6 mice bearing transplanted KPROR1 lung tumors. N=3-4 mice per group. One-way ANOVA with Tukey's post-test. B, C) Inhibitory receptor expression (B) and cytokine production upon ex vivo restimulation with PMA and ionomycin (C) in CD8+GFP+ CAR-T cells in lungs 8 days post-infusion into B6 mice bearing transplanted KP-ROR1 lung tumors. N=3-4 mice per group. One-way ANOVA with Tukey's post-test. D) Survival of B6 mice bearing transplanted KP-ROR1 tumors and treated with the indicated T cell products. N=4 mice per group. Log-rank Mantel-Cox test. Data are representative of 2 independent experiments.

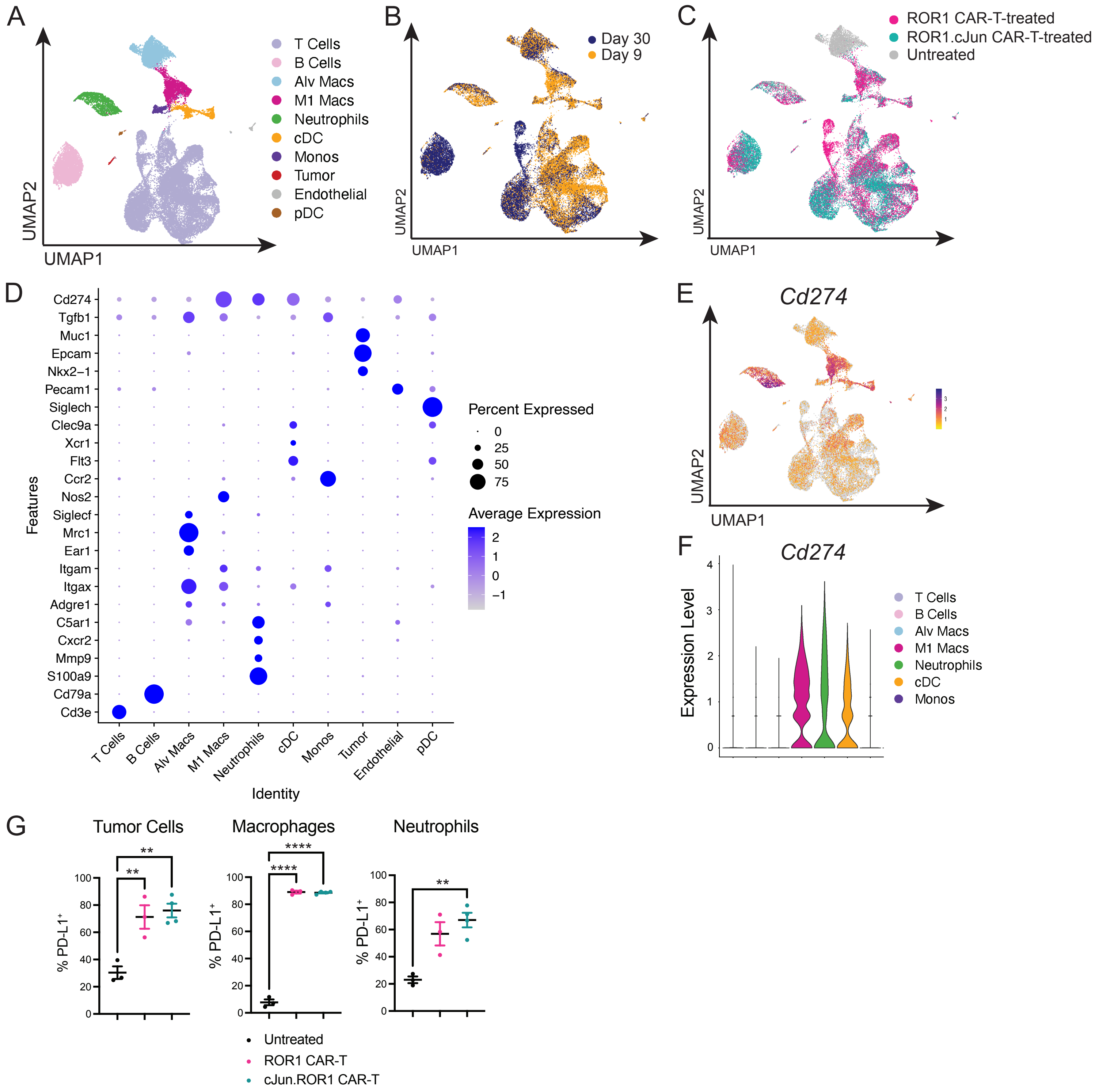


**Figure S5. Annotation of TME single cell RNAseq clusters.** A) Unsupervised clustering of live cells sorted from KP$^{ROR1}$ tumors of mice left untreated, or 9 or 30 days post-infusion with ROR1 CAR-T or cJun.ROR1 CAR-T cells. B, C) Clusters in (A) colored by time point (B) or treatment group (C). D) Dot plot showing expression of various lineage-defining genes across clusters in (A). E,F) UMAP (E) and violin plot (F) showing expression of *Cd274* (encodes PD-L1) across clusters in (A). G) Expression of PD-L1 on CD45-EpCAM+ROR1+ KP tumor cells, CD11c+F4/80+ macrophages, and CD11b+Ly6G+ neutrophils within KP$^{ROR1}$ lung tumors left untreated (black) or 9 days post-infusion of ROR1 CAR-Ts (pink) or cJun.ROR1 CAR-Ts (teal). N=3-5 mice per group. One-way ANOVA with Tukey's post-test.

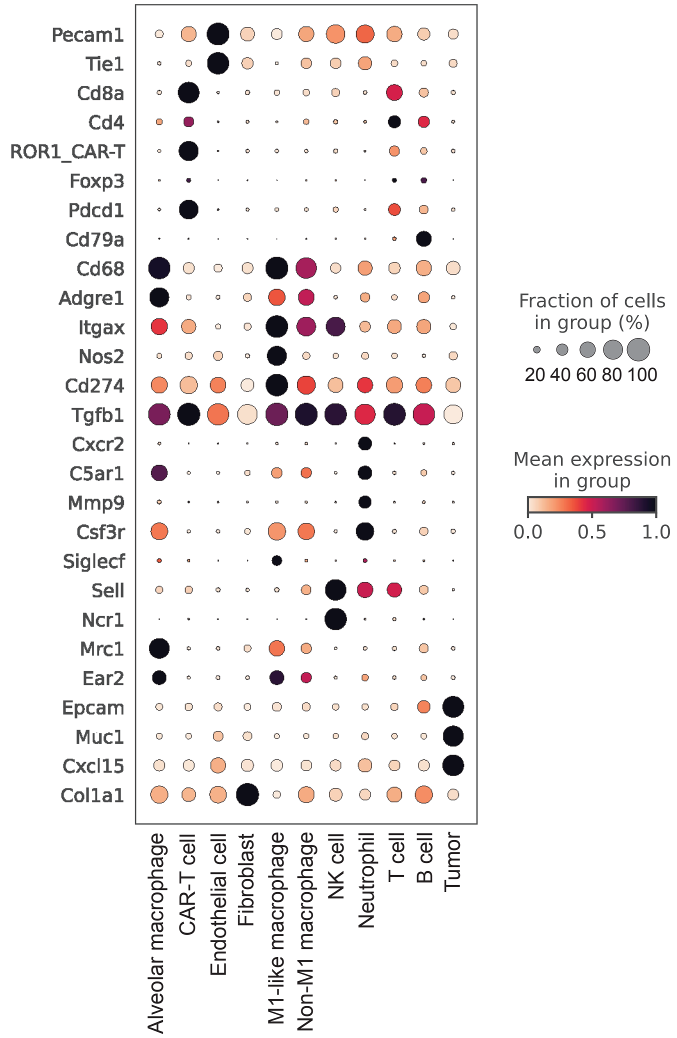


**Figure S6. Annotation of 10X Xenium clusters.** Lungs from $KP^{ROR1}$ mice were harvested 7 days post-treatment with ROR1 CAR-Ts + vehicle, ROR1 CAR-Ts + anti-PD-L1, cJun.ROR1 CAR-Ts + vehicle, or cJun.ROR1 CAR-Ts + anti-PD-L1 and analyzed by 10X Xenium. Dot plot summarizes expression of various genes defining each cluster.
